# Interferon-Inducible Guanylate-Binding Protein 5 Inhibits Replication of Multiple Viruses by Binding to the Oligosaccharyltransferase Complex and Inhibiting Glycoprotein Maturation

**DOI:** 10.1101/2024.05.01.591800

**Authors:** Shaobo Wang, Wanyu Li, Lingling Wang, Shashi Kant Tiwari, William Bray, Lujing Wu, Na Li, Hui Hui, Alex E. Clark, Qiong Zhang, Lingzhi Zhang, Aaron F. Carlin, Tariq M. Rana

## Abstract

Viral infection induces production of type I interferons and expression of interferon-stimulated genes (ISGs) that play key roles in inhibiting viral infection. Here, we show that the ISG guanylate-binding protein 5 (GBP5) inhibits N-linked glycosylation of key proteins in multiple viruses, including SARS-CoV-2 spike protein. GBP5 binds to accessory subunits of the host oligosaccharyltransferase (OST) complex and blocks its interaction with the spike protein, which results in misfolding and retention of spike protein in the endoplasmic reticulum likely due to decreased *N*-glycan transfer, and reduces the assembly and release of infectious virions. Consistent with these observations, pharmacological inhibition of the OST complex with NGI-1 potently inhibits glycosylation of other viral proteins, including MERS-CoV spike protein, HIV-1 gp160, and IAV hemagglutinin, and prevents the production of infectious virions. Our results identify a novel strategy by which ISGs restrict virus infection and provide a rationale for targeting glycosylation as a broad antiviral therapeutic strategy.

**Highlights:**

1. The interferon-stimulated gene GBP5 is induced by SARS-CoV-2 infection in vitro and in vivo.
2. ER-localized GBP5 restricts N-linked glycosylation of SARS-CoV-2 spike protein, leading to protein misfolding and preventing transport to the Golgi apparatus.
3. GBP5 binds to OST complex accessory proteins and potentially blocks access of the catalytic subunit to the spike protein.
4. GBP5 inhibits N-glycosylation of key proteins in multiple viruses, including SARS-CoV-2
5. Pharmacological inhibition of OST blocks host cell infection by SARS-CoV-2, variants of concern, HIV-1, and IAV.

**Significance:** Viral infection induces production of type I interferons and expression of interferon-stimulated genes (ISGs) that play key roles in inhibiting viral infection. We found that the interferon-stimulated gene GBP5 is induced by SARS-CoV-2 infection in vitro and in vivo. GBP5 inhibits N-glycosylation of key proteins in multiple viruses, including SARS-CoV-2. Importantly, pharmacological inhibition of Oligosaccharyltransferase (OST) Complex blocks host cell infection by SARS-CoV-2, variants of concern, HIV-1, and IAV, indicating future translational application of our findings.

## Introduction

Since its emergence in December 2019, severe acute respiratory syndrome coronavirus 2 (SARS-CoV-2), the causative agent of coronavirus disease 2019 (COVID-19), has continued to pose a global public health threat. As of June 2022, more than 540 million cases of SARS-CoV-2 infection have been documented worldwide and have caused more than 6 million deaths (WHO COVID-19 Dashboard)^1^. Although several effective vaccines have been deployed worldwide, waves of new infections have followed the emergence of SARS-CoV-2 variants of concern (VOCs)^2–5^. Several of these VOCs, especially B.1.1.529 (Omicron), exhibit increased resistance to convalescent sera, vaccinee sera, and monoclonal antibodies that have been authorized for clinical use^3,6^, suggesting a need for vaccine refinement as well as the development of new approaches to COVID-19 treatment.

Viral infection induces activation of the innate immune system and the production of antiviral cytokines and other mediators. Among these, the type I interferons (e.g., IFN-α, IFN-β) potently induce expression of a panel of host interferon-stimulated genes (ISGs). Expression of ISGs gives rise to a localized antiviral state that is instrumental in the control and clearance of virus^7^. The importance of type I IFNs in combating viral infection is illustrated by the identification of genetic defects in IFN regulatory factors and the presence of neutralizing antibodies against type I IFNs as two risk factors for developing severe COVID-19 pneumonia^8,9^. In addition, upper airway epithelial cells from children have been shown to express elevated basal levels of pattern recognition receptors and to induce ISGs faster and more robustly compared with the corresponding cells from adults, providing a possible explanation for the lower SARS-CoV-2 infection rates and reduced risk of severe COVID-19 in children compared with adults^10–12^. Although IFNs have been considered possible therapeutic candidates for the treatment of COVID-19, experience to date indicates that they can promote local and systemic inflammation and can exacerbate viral disease, including COVID-19^13,14^. Therefore, understanding the mechanism of action of individual ISGs in the viral life cycle might inform the development of more specific antiviral therapeutic strategies with fewer undesirable side effects.

ISGs function through diverse mechanisms that interfere with all stages of the viral life cycle. We previously demonstrated that the ISG cholesterol 25-hydroxylase (CH25H) inhibits SARS-CoV-2 host cell entry by depleting accessible cholesterol at the plasma membrane, which is necessary for spike (S) protein-mediated fusion independently of viral fusogenic proteins^15^. Other ISGs also act at specific steps in viral replication and release; for example, OAS1 and BST-2 restrict viral RNA replication and virion release, respectively^15–19^. Notably, ISGs that function in virion assembly and release are generally less well-characterized than ISGs that function in earlier stages of the viral life cycle^20^.

Glycosylation is one of the most common and important post-translational modifications of host and viral proteins, and it plays key roles in determining protein structure, stability, and function^21,22^. Glycoproteins undergo either O-linked glycosylation at serine or threonine residues and/or N-linked glycosylation at asparagine residues within the endoplasmic reticulum (ER). In the case of N-linked glycosylation, a 14-sugar precursor structure, Glc_3_Man_9_GlcNAc_2_ (Glc, glucose; Man, mannose; GlcNAc, N-acetylglucosamine), attaches to a long-chain polyisoprenol dolichol phosphate that serves as the glycan donor in the ER^21^. The donor chain is then transferred to an asparagine (Asn, N) residue in the sequon Asn-X-serine/threonine (where X is any amino acid other than proline) in the substrate protein by the oligosaccharyltransferase (OST) complex, which consists of STT3A and STT3B catalytic subunits and various accessory subunits including RPN1, RPN2, and DDOST^23^. Glycosylated proteins are further processed in the ER and then, for glycoproteins targeted to the secretory pathway, in the Golgi apparatus. The glucose and mannose moieties are sequentially removed during assisted protein folding, ER quality control, and trafficking. Properly folded proteins that are targeted to the secretory pathway then traffic to the Golgi apparatus where they acquire hybrid and complex carbohydrate branches^21^. Although many enveloped human viruses hijack host glycosylation machineries for viral glycoprotein maturation, virion assembly, epitope shielding, and sometimes even for noncanonical structural support^24^, it is not currently known whether ISGs induced in the host cell are able to interfere with such a ubiquitous viral dependency pathway.

Here, we report a novel mechanism of antiviral activity by ISGs that centers on inhibition of viral protein glycosylation. Using SARS-CoV-2 S protein as a model viral glycoprotein, we show that the ISG guanylate-binding protein 5 (GBP5), a member of the large family of IFN-inducible GTPases that play crucial roles in innate immunity, interferes with OST-mediated N-linked glycosylation of S protein, resulting in its misfolding and retention in the ER, and reducing SARS-CoV-2 virion assembly and release. Notably, we demonstrate that pharmacological inhibition of the OST also blocks protein glycosylation and virion production by additional viruses, including influenza A virus (IAV), human immunodeficiency virus (HIV), Middle East respiratory syndrome coronavirus (MERS-CoV), and SARS-CoV-2 VOCs B.1.617.2 (Delta), B.1.351 (Beta), and B.1.1.529 (Omicron), adding to existing evidence of the antiviral effects of OST inhibition^25,26^. To the best of our knowledge, this is the first study to describe a mechanism by which ISGs regulate viral protein glycosylation and may represent a new strategy for the development of broad-spectrum antiviral therapies.

## Results

### GBP5 is induced by SARS-CoV-2 infection and inhibits viral assembly and release

To investigate the potential mechanisms by which ISGs may interfere with viral infection at later points in the life cycle, we analyzed published RNA-seq data from SARS-CoV-2-infected Calu-3 and A549-ACE2 cell lines, both of which are human lung adenocarcinoma cells and the latter of which overexpresses the cellular receptor for SARS-CoV-2, ACE2^27^. As expected, many ISGs were upregulated by SARS-CoV-2 infection, including multiple members of the GBP family (Fig 1a and S1a-e). We observed similar induction of GBP transcripts in A549 cells infected by the respiratory viruses human parainfluenza virus type 3 (HPIV3) and respiratory syncytial virus (RSV) (Fig 1a). Notably, these genes were not upregulated in A549 cells infected by IAV, which is consistent with the known ability of IAV infection to inhibit the IFN signaling pathway (Fig 1a). GBP genes were also upregulated in SARS-CoV-2 infected human alveolar epithelial type-2 (AT2) organoids and lung tissues from COVID-19 patients, with GBP5 being highly upregulated in both cases (∼44-fold and ∼16-fold vs mock-infected cells and normal lung tissue, respectively; Fig 1b,c)^27,28^. Intriguingly, analysis of scRNA-seq datasets from SARS-CoV-2-susceptible upper airway cells revealed higher expression of GBP5 in cells from healthy children vs healthy adults, and a larger induction of GBP5 in cells from SARS-CoV-2-infected children vs infected adults (Fig 1d and S1f), which is consistent with the recent suggestion that higher basal activity of the innate immune system in children may explain their lower SARS-CoV-2 infection rate and decreased risk of developing COVID-19 compared with adults^10^. We further confirmed the induction of GBP5 expression by SARS-CoV-2 USA-WA1/2020 (referred to as wild-type [WT] strain) in Calu-3 cells (Fig S1g-S1h).

**Figure 1.**
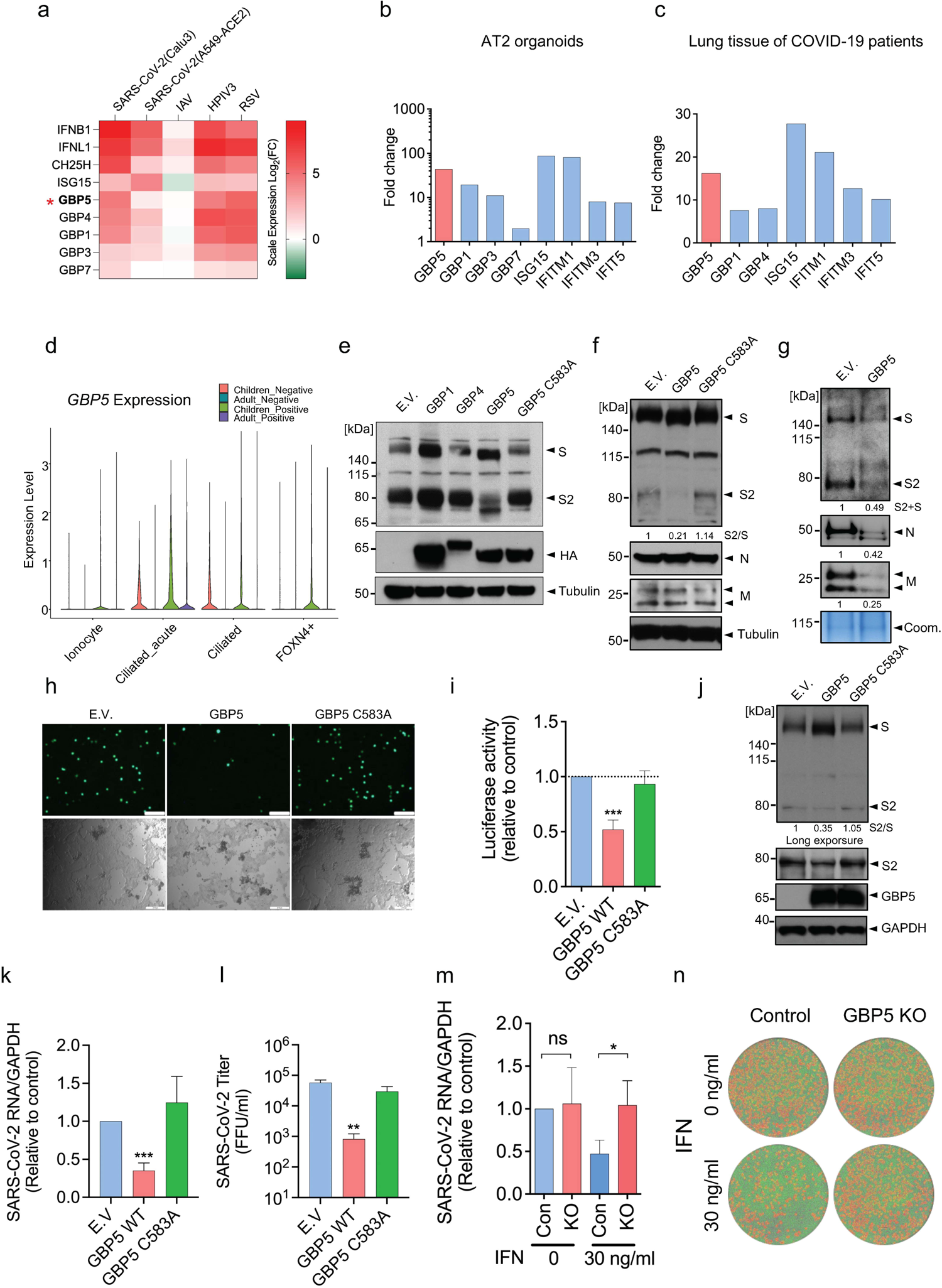
GBP5 expression is induced in lung cells and organoids by SARS-CoV-2 infection and restricts virus release. a. GBPs and other ISGs are induced in SARS-CoV-2-infected human lung epithelial cell lines. RNA-seq analysis of ISGs in Calu-3 and A549-ACE2 cells infected with SARS-CoV-2 USA-WA1/2020 for 24 h at an MOI of 2; in A549 cells infected with IAV at an MOI of 5 for 9 h; and in A549 cells infected with HPIV3 and RSV at an MOI of 2 for 24 h. Log2 fold change in gene expression is shown relative to mock-infected cells. Data are taken from reference ^27^. b, c. GBP5 and other ISGs are induced in SARS-CoV-2-infected human alveolar type II cell organoids and in lung tissue from COVID-19 patients. RNA-seq analysis of (b) AT2 organoids infected with SARS-CoV-2 at an MOI of 1 for 48 h or (c) lung tissues from two COVID-19 patients and two healthy donors. Fold change in gene expression is shown relative to mock-infected organoids (b) or lung tissue from healthy donors (c). Data are taken from reference ^27^. d. GBP5 is induced at higher levels in upper airway cells from SARS-CoV-2-infected children compared with adults. scRNA-seq analysis of GBP5 expression in nasal samples taken from healthy (negative) or SARS-CoV-2-infected (positive) children and adults (Negative children n=18, positive children n=24, negative adults n=23 and positive adults n=21). Violin plots show the relative expression of GBP5 in the indicated upper airway cell types. Data are taken from reference ^10^. e. GBP5 suppresses S protein cleavage. Western blot analysis of 293T cells transfected for 48 h with empty vector (EV) or the indicated HA-tagged GBP family proteins and co-transfected with SARS-CoV-2 S protein. Blots were probed with an antibody that recognizes the cleaved S2 subunit and full-length S protein or with anti-HA antibody. Tubulin was probed as a loading control. f, g. GBP5 inhibits the release of SARS-CoV-2 VLPs. Western blot analysis of whole cell lysates (f) or concentrated supernatants (g) of 293T cells transfected for 48 h with VLPs encoding SARS-CoV-2 S protein (S), nucleocapsid (N), and membrane (M) proteins and co-transfected with HA-tagged GBP5, HA-GBP5-C583A, or EV. Blots were probed with antibodies against the indicated viral proteins or tubulin. Coomassie-stained gels served as a loading control for VLP-containing supernatants (arrows). h, i. GBP5 overexpression inhibits the release of SARS-CoV-2 pseudovirus. EGFP-expressing (h) or luciferase-expressing (i) SARS-CoV-2 pseudoviruses were produced in 293T cells transfected with EV, GBP5, or GBP5-C583A. After 24 h, supernatants were collected and used to infect Calu-3 cells for 2 h. Infectivity of Calu-3 cells was analyzed by fluorescence microscopy (h) or luciferase assay (i). Mean ± SD of n = 3. ***p < 0.001 by Student’s t-test. Scale bar, 100 μm. j-l. GBP5 suppresses infection by live SARS-CoV-2. 293T/ACE2/TMPRSS2 cells were transfected with EV, GBP5, or GBP5-C583A and infected with SARS-CoV-2 USA-WA1/2020 strain at a MOI of 0.1. After 24 h, cells and supernatants were collected and analyzed. (j) Western blot analysis of intact S protein, cleaved S2 subunit, GBP5, and GAPDH (loading control). RT-qPCR analysis of viral RNA in cell extracts (k) and viral titer in the supernatant assessed by FFU (l). Mean ± SD of n = 3. *p < 0.05, **p < 0.01 by Student’s t-test. m-n. GBP5 contributes to the antiviral activity of IFN-γ against SARS-CoV-2 in lung epithelium. GBP5 KO or control Calu3 cells were pretreated with 30 ng/ml IFN-γ for 16 h and infected with SARS-CoV-2 USA-WA1/2020 strain at a MOI of 0.1. After 48h, cells and supernatants were collected and analyzed. The viral RNA in cell extracts was quantified by RT-qPCR (m) and viral titer was evaluated by FFU assay (n). Mean ± SD of n = 3. *p < 0.05, **p < 0.01 by Student’s t-test.

GBP5 has been proposed to inhibit cleavage of viral glycoproteins by the Golgi-resident protease furin^29^, which is also responsible for cleavage of the ∼180 kDa SARS-CoV-2 S protein into subunits S1 (∼100 kDa) and S2 (∼80 kDa)^25^. Therefore, we analyzed production of the S2 subunit in 293T cells co-transfected with SARS-CoV-2 S protein and GBP5, GBP1, or GBP4. Indeed, overexpression of GBP5, but not GBP4 or GBP1, not only inhibited S protein cleavage and the appearance of S2 but also decreased the apparent molecular weight (MWa) of the S protein (Fig 1e), suggesting that GBP5 overexpression had additionally influenced an unknown modification of the full-length S protein. Post-translational prenylation of GBP5 at cysteine 583 allows GBP5 to associate with membranes, which is crucial for its ability to inhibit viral glycoprotein cleavage ^30^. As expected, we observed no change in S2 production or in the S protein MWa in 293T cells overexpressing GBP5 harboring a cysteine to alanine mutation (C583A) (Fig 1e), indicating that the ability of GBP5 to inhibit S protein cleavage and decrease S protein MWa requires its membrane anchor motif.

To evaluate the effect of GBP5 and GBP5-C583A overexpression on SARS-CoV-2 virion assembly, we constructed replication-incompetent virus-like particles (VLPs) bearing SARS-CoV-2 nucleocapsid (N), membrane (M), and S proteins in control or GBP5-overexpressing 293T cells^31,32^ and examined N, M, and S protein levels both intracellularly and in the supernatant as a measure of VLP secretion. Cells co-transfected with VLPs and GBP5 showed reduced S2 production and a reduction in MWa that was not observed in GBP5-C583A-expressing cells, while intracellular levels of N and M proteins were not affected (Fig 1f). However, S, N, and M protein levels in the supernatants were all strikingly reduced by GBP5 overexpression, suggesting a marked inhibition of VLP assembly and release (Fig 1g). To test this further, we produced EGFP- or luciferase-expressing VSV pseudoviruses bearing SARS-CoV-2 S protein in control, GBP5-, or GBP5-C583A-overexpressing 293T cells, collected the pseudovirus-containing supernatant, and measured pseudovirus titer by addition of the supernatants to Calu-3 cells. Analysis of Calu-3 cells by fluorescence microscopy (Fig 1h) or by luciferase assays (Fig 1i) showed that GBP5 overexpression significantly reduced pseudovirus infectivity compared with control and GBP5-C583A-overexpressing cells (Fig 1h,i).

To confirm these findings in the context of authentic virus infection, we examined the effect of GBP5 or GBP5-C583A overexpression on S protein processing and infection with SARS-CoV-2 in HEK 293T cells expressing the SARS-CoV-2 co-receptors ACE2 and TMPRSS2. We again observed inhibition of S2 production and the alteration in S protein MWa in infected cells overexpressing GBP5, but not GBP5-C583A (Fig 1j), as well as reduced viral RNA levels in infected cells (Fig 1k). Importantly, culture supernatants of SARS-CoV-2 infected GBP5-overexpressing cells contain significantly less infectious virions than supernatants of control or GBP5-C583A-overexpressing cells (Fig 1l), indicating effective suppression of virion release. Taken together, these results show that GBP5 inhibits SARS-CoV-2 virion assembly and release by interfering with S protein maturation.

We next determined the importance of GBP5 in the context of an interferon response. We confirmed that GBP5 expression is inducible by IFN-γ, and that pre-treatment with IFN-γ inhibits SARS-CoV-2 infection in Calu-3 cells (Fig S1h-S1i). To determine whether GBP5 plays a role in the restriction of viral replication by IFN-γ, we genetically depleted Calu-3 cells of GBP5 by CRISPR-Cas9 (Fig S1j). Control and GBP5 KO Calu-3 cells were pre-treated with IFN-γ and challenged with SARS-CoV-2. We observed that IFN-γ suppression of SARS-CoV-2 replication was significantly reversed by GBP5 KO, as indicated by the rescue of intracellular and supernatant viral RNA levels (Fig 1m and S1k). Furthermore, GBP5 KO reduced infectious virion titer in the culture supernatant (Fig 1n). Finally, we tested the effects of GBP5 depletion by shRNA on SARS-CoV-2 infection of primary human bronchial epithelial cells (Fig S1l). We found that GBP5 depletion increased SARS-CoV-2 replication in these primary cells with IFN-γ treatment (Fig S1m). Collectively, these data provide evidence that GBP5 is an important contributor to the antiviral response induced by IFN-γ.

### GBP5 alters N-linked glycosylation and trafficking of SARS-CoV-2 S protein

To understand the mechanism by which GBP5 overexpression could not only decrease the MWa of SARS-CoV-2 S protein but also affect its proteolytic processing, we focused on the potential involvement of N-linked glycosylation for two main reasons: first, addition or removal of large carbohydrate complexes might potentially explain the observed changes in S protein MWa, and second, correct N-linked glycosylation is essential for trafficking of glycoproteins, including viral S proteins, along the secretory pathway to the Golgi compartment, where they could be processed by proprotein convertases such as furin^33–35^. Additionally, inhibition of N-linked glycosylation has been shown to result in retention of glycoproteins in the ER^36,37^. Thus, we hypothesized that GBP5 may affect S protein processing by inhibiting its glycosylation, resulting in its retention in the ER, and inhibition of cleavage.

To examine this, we first determined whether removal of N-linked glycosylation eliminates the reduction in S protein MWa observed upon GBP5 overexpression. To this end, we incubated lysates from SARS-CoV-2 S protein-transfected control, GBP5-, or GBP5-C583A-overexpressing 293T cells with endoglycosidase H (Endo H) or peptide N-glycosidase F (PNGase F), which cleave high mannose-type N-linked glycans and all N-linked glycans, respectively (Fig 2a). The activity of PNGase F, therefore, allows us to examine whether the decrease in the MWa of S protein under GBP5 could survive the disappearance of N-linked glycans and any alteration therein. Spike-containing lysates from control cells, or cells overexpressing GBP5 or GBP5 C583A, were digested by these two enzymes. When compared with mock treated sample, our results showed that with N-linked glycans removed by PNGase F, electrophoretic mobility of bands corresponded to unglycosylated S and S2 proteins (Fig 2b, lanes 1-3 and 7-9), confirming that the observed change in MWa is indeed due to decreased N-linked glycosylation.

**Figure 2.**
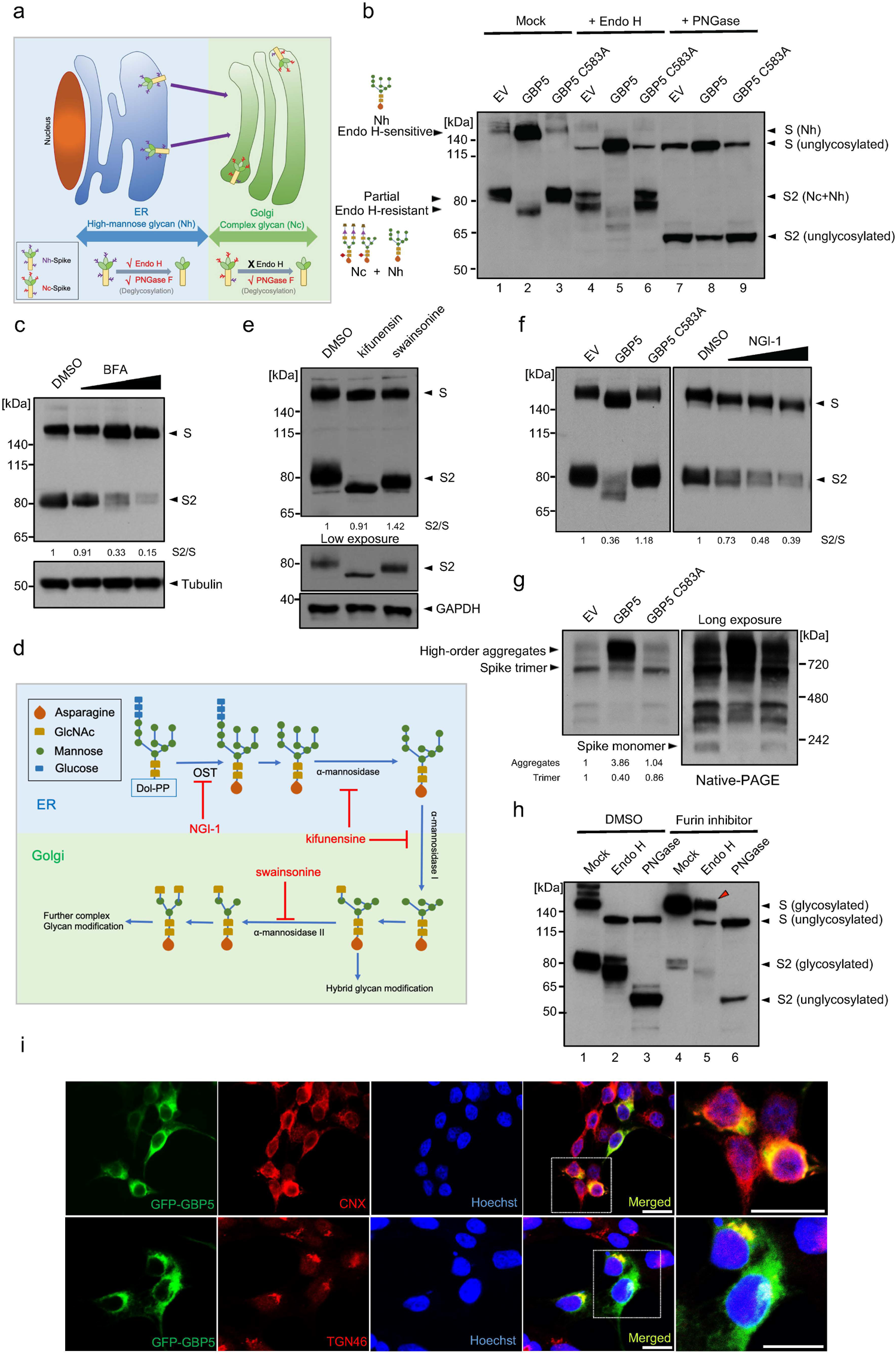
GBP5 suppresses SARS-CoV-2 S protein glycosylation and trafficking. a. Schematic showing the relationship between Endo H and PNGase F sensitivity and protein localization along the secretory pathway. Glycoproteins in the ER harbor high mannose-type (Nh) glycans that are sensitive to both Endo H and PNGase F digestion. Glycoproteins in the Golgi apparatus harbor complex type (Nc) glycans that are sensitive to PNGase F but not to Endo H digestion. b. Sensitivity of SARS-CoV-2 S protein to Endo H and PNGase F digestion. Western blot analysis of glycosylated and unglycosylated S protein in 293T cells transfected with empty vector (EV), GBP5, or GBP5-C583A. Cell lysates were digested for 1 h with Endo H or PNGase F. Blots were probed with anti-S2 subunit antibody. c. Brefeldin A (BFA) inhibits SARS-CoV-2 S protein proteolytic cleavage. Western blot analysis of lysates from 293T cells transfected for 42 h with SARS-CoV-2 S protein and then treated with DMSO or 0.012, 0.05, or 0.2 μM BFA for 6 h. Blots were probed with antibodies against S protein and tubulin (loading control). d. Schematic showing sequential N-linked glycosylation through the secretory pathway. Only enzymes inhibited by NGI-1, kifunensine, and swainsonine are shown. The OST complex transfers glycans from dolichol to arginine residues in the substrate protein; α-mannosidase I in the ER and Golgi apparatus trims mannose moieties to allow for hybrid type modification; and α-mannosidase II in the Golgi further trims mannose to commit the protein for complex type modification. e. Inhibition of α-mannosidase I and II do not block SARS-CoV-2 S protein cleavage. Western blot analysis of 293T cells transfected for 4 h with SARS-CoV-2 S protein and then incubated with DMSO, 25 μM kifunensine, or 40 μM swainsonine for an additional 48 h. Blots were probed with antibodies against S protein or GAPDH (loading control). f. Inhibition of OST inhibits SARS-CoV-2 S protein cleavage and prevents N-glycosylation. Western blot analysis of 293T cells transfected for 42 h with SARS-CoV-2 S protein and then incubated for 6 h with DMSO or 1, 2, or 4 μM NGI-1. Blots were probed with anti-S protein antibody. g. GBP5 induces misfolding of S protein. 293T cells were transfected for 48 h with EV, GBP5, or GBP5-C583A and subjected to native PAGE. Blots were probed with anti-S protein antibody. h. Inhibition of furin leads to accumulation of Endo H-resistant full-length S protein. Western blot analysis of 293T cells transfected for 4 h with SARS-CoV-2 S protein and then treated with DMSO or 100 μM CMK for 48 h. Lysates were digested with Endo H or PNGase F for 1 h. Blots were probed with anti-S protein antibody. i. GBP5 is present in the ER but not the Golgi apparatus. Fluorescence microscopy of 293T cells transfected with EGFP-GBP5 and stained with antibodies against the ER marker calnexin (CNX, upper row) or the trans-Golgi marker TGN46 (lower row). Nuclei were stained with Hoechst 33342. Scale bar, 50 μm.

Next, based on N-linked glycan composition on S protein, we analyzed changes in S protein localization. Initial N-linked glycans added onto glycoproteins in the ER are high mannose-type, which is sensitive to Endo H digestion. As glycoproteins traffic along the secretory pathway, high mannose-type glycans are modified into Endo H-resistant, hybrid and complex glycans by enzyme cascades in Golgi compartments (Fig 2a)^21^. Accordingly, glycoproteins could only bear complex N-linked glycans and resist Endo H digestion after having reached the Golgi apparatus^37–40^.

Western blot analysis showed full-length S protein was Endo H-sensitive, while cleaved S protein (S2) was partially Endo H-resistant (Fig 2b, lanes 1-3 and 4-6). This indicates that cleaved S protein harbored complex-type N-linked glycans, which suggest that S protein molecules that were cleaved must have trafficked through Golgi compartments. On the other hand, full-length S protein only harbored high mannose-type N-linked glycans, which suggest that uncleaved S protein molecules were localized in the ER and had yet to traffic to the Golgi apparatus where it could obtain complex glycan modification. Importantly, overexpression of GBP5 led to an accumulation of full-length S protein, which localized to the ER, and a depletion of cleaved S protein (S2), which had trafficked through the Golgi apparatus, compared to transfection of an empty vector or GBP5-C583A (Fig 2b, lanes 4-6). In other words, GBP5 overexpression retained S protein in the ER compared to control or GBP5-C583A overexpression. Finally, we observed that Endo H removed the difference in the MWa of full-length S protein but not of S2 (Fig 2b, lanes 1-3 and 4-6), suggesting that alteration in N-linked glycosylation under GBP5 overexpression took place when N-linked glycans on S protein were still high mannose-type, likely when S protein molecules were still in the ER. Collectively, these observations suggest that GBP5 reduces S protein N-linked glycosylation in the ER and decreases S protein trafficking to Golgi compartments where furin-catalyzed proteolytic processing occurs.

Our hypothesis implies that a reduction in trafficking prevents S protein cleavage. To confirm this proposition, we treated S protein-expressing cells with brefeldin A (BFA), an inhibitor of transport from the ER to the Golgi apparatus. This resulted in a dose-dependent inhibition of S protein cleavage (Fig 2c), verifying that blocking ER-to-Golgi trafficking impedes S protein proteolytic processing.

To identify the precise step in glycan maturation at which GBP5 acts, we asked whether inhibition of select enzymes in the N-linked glycosylation sequence that function prior to complex modification would mimic GBP5 overexpression. SARS-CoV-2 S protein-transfected HEK 293T cells were incubated with kifunensine, an inhibitor of α-mannosidase I in the ER and Golgi; swainsonine, an inhibitor of Golgi α-mannosidase II; and NGI-1, an inhibitor of the OST complex in the ER^36,41^ (Fig 2d). Kifunensine and swainsonine did not induce a shift in the MWa of full-length S protein or decrease its proteolytic processing at nontoxic concentrations, although the MWa of S2 was reduced (Fig 2e, Fig S2A). However, NGI-1 dose-dependently decreased S protein cleavage and concomitantly reduced the S protein MWa (Fig 2f, S2c), recapitulating major effects of GBP5 overexpression on S protein processing. These results therefore suggest that GBP5 may impede S protein processing by interfering with N-glycan transfer onto S protein by the OST complex function in the ER.

Inadequate N-linked glycosylation often leads to protein misfolding, which prevents trafficking from the ER to the Golgi^42,43^. Indeed, when lysates of SARS-CoV-2 S protein-transfected HEK 293T cells were analyzed by native PAGE, substantial accumulation of higher-order S protein aggregates was observed in cells overexpressing GBP5, but not in control or GBP5-C583A-overexpressing cells, and the abundance of monomeric and trimeric S protein decreased in parallel (Fig 2g). Thus, GBP5 overexpression promotes S protein misfolding and aggregation, consistent with importance of proper glycosylation in protein folding and trafficking.

Previously, GBP5 has been proposed to inhibit the proteolytic maturation of various viral glycoproteins by interfering with furin activity^29^. However, our hypothesis and results so far entail that the inhibition of a specific viral glycoprotein proteolysis is mainly due to retention in the ER, and not decreased furin activity. If furin activity is inhibited alone in GBP5-overexpressing cells without effects on glycoprotein trafficking, we would likely see full-length S protein that acquired complex glycans in the Golgi apparatus but could not be proteolytically processed. Indeed, treatment with a furin inhibitor, Decanoyl-RVKR-CMK (CMK), potently inhibited proteolytic processing of S protein and simultaneously gave rise to a strong signal of Endo H-resistant full-length S protein (Fig 2h, Fig S2d). On the other hand, full-length S protein was completely sensitive to Endo H in GBP5-overexpressing cells (Fig 2b, lane 5). These observations suggest that inhibition of furin alone does not prevent S protein trafficking to the Golgi apparatus as GBP5 does, and show that the effects of GBP5 on S protein maturation are not the results of altered furin activity.

Another question that arises from our hypothesis is that GBP5 has only been reported to localize to the Golgi apparatus. If GBP5 functions no later than glycan transfer, then it may need to localize to the ER where the OST complex and the enzymes that synthesize dolichol-linked glycan precursors localize^21^. To determine whether it is possible for GBP5 to target ER processes, we performed immunofluorescence staining. We found that GFP-tagged GBP5 co-localized partially with both the ER marker calnexin and the Golgi marker TGN46 (Fig 2i). To exclude the possibility of mislocalization due to ectopic expression, we then examined endogenous GBP5 localization in HEK 293T cells treated with IFN-γ to induce GBP5 expression. Once again, we observed GBP5 partially co-localized with calnexin (Fig. S2e). These results indicate that GBP5 is indeed found in the ER and may interact with ER resident proteins involved in glycan synthesis and transfer. Several other reports have also shown that GBP5 is diffusely distributed in the cytoplasm, characteristic of ER distribution, while only partially co-localizing with Golgi markers^29,44,45^. Overall, our results favor a model in which GBP5 interferes with N-linked glycosylation of SARS-CoV-2 S protein in the ER, reduces its trafficking to post-Golgi compartments and thereby inhibits its proteolytic maturation.

### GBP5 interacts with the OST complex via the non-catalytic accessory proteins

As noted earlier (Fig 2f), inhibition of the OST complex with NGI-1 largely mimicked the GBP5-induced reduction in cleavage and MWa of SARS-CoV-2 S protein, suggesting that GBP5 might act by interfering with OST-mediated S protein glycosylation. To test this, we expressed FLAG-tagged GBP5 and MYC-tagged RPN2, RPN1, and DDOST in HEK 293T cells, immunoprecipitated lysates with anti-FLAG antibody and probed for co-immunoprecipitation of OST subunits by anti-MYC western blotting. Indeed, all three accessory subunits were shown to co-immunoprecipitate with FLAG-GBP5 (Fig 3a–c). Moreover, immunoprecipitation of FLAG-GBP5, but not FLAG-GBP5-C583A, pulled down endogenous RPN2, RPN1, and DDOST in HEK 293T cells (Fig 3d,e). To determine whether these interactions occur under endogenous expression levels of GBP5 and the OST subunits, we treated HEK 293T cells with IFN-γ to induce GBP5 expression, and immunoprecipitated GBP5 from the cell lysate using an anti-GBP5 antibody. All of RPN1, RPN2, and DDOST were detected in the immunoprecipitates by western blot (Fig S3a).

**Figure 3.**
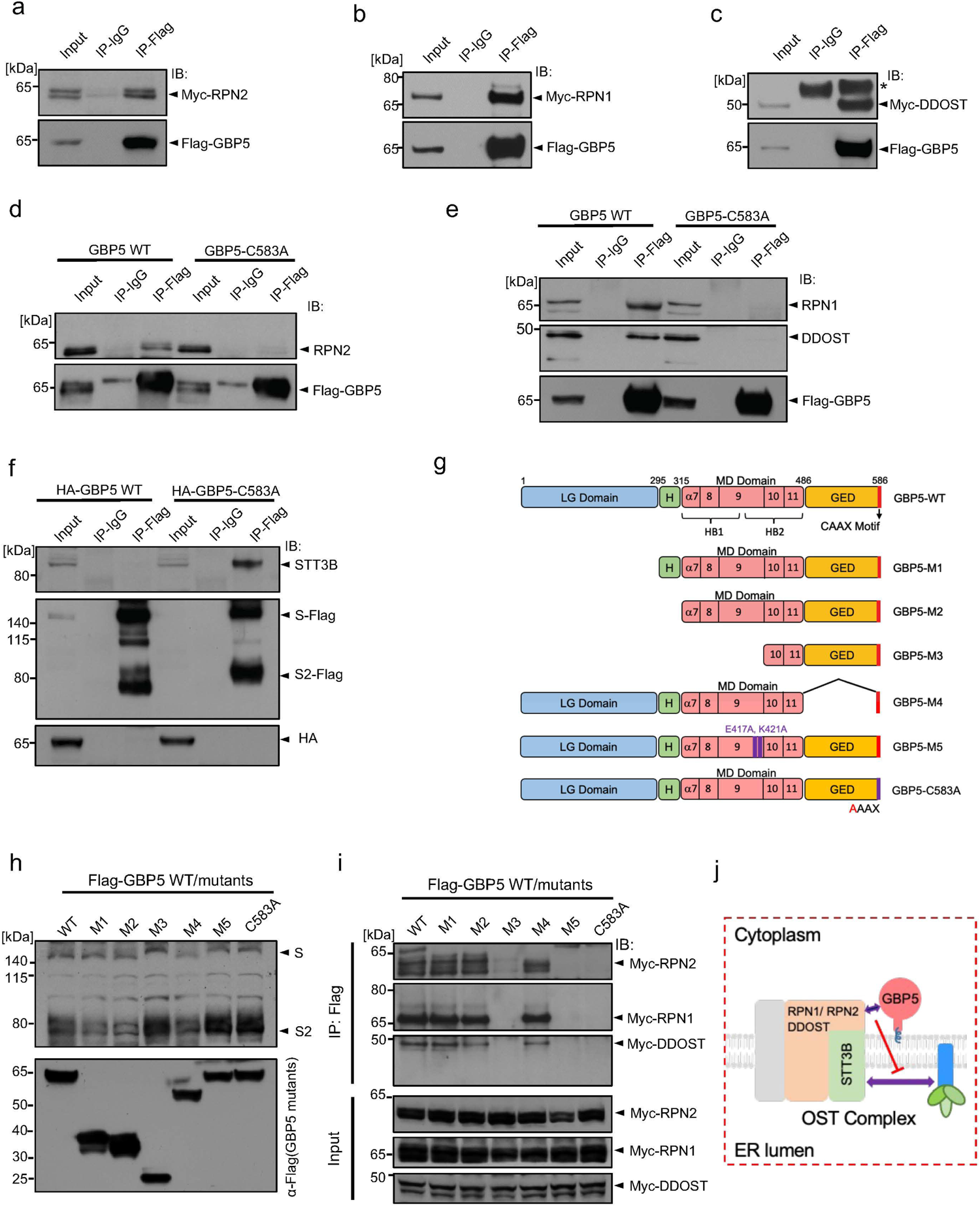
GBP5 interacts with the OST complex and blocks its interaction with SARS-CoV-2 S protein. a-c. Ectopically expressed FLAG-tagged GBP5 interacts with ectopically expressed OST subunits. Western blot analysis of 293T cells transfected for 48 h with FLAG-tagged GBP5 and either MYC-RPN2 (A), MYC-RPN1 (B), or MYC-DDOST (C). Lysates were immunoprecipitated with anti-FLAG antibody and blots were probed with anti-MYC or anti-FLAG antibodies. d, e. Ectopically expressed GBP5 interacts with endogenous OST subunits. Western blot analysis of 293T cells transfected for 48 h with FLAG-tagged GBP5 or GBP5-C583A. Lysates were immunoprecipitated with anti-FLAG antibody and blots were probed with antibodies against FLAG and RPN2 (D) or RPN1 and DDOST (E). f. GBP5 blocks SARS-CoV-2 S protein binding to endogenous STT3B. Western blot analysis of 293T cells transfected for 48 h with plasmids encoding FLAG-tagged S protein and HA-tagged GBP5 or GBP5-C583A. Lysates were immunoprecipitated with anti-FLAG antibody and blots were probed with antibodies against STT3B, FLAG, or HA. g. Schematic showing mutant constructs of GBP5. M1: deletion of the large GTPase (LG) domain; M2: deletion of the LG domain and the hinge region connecting the LG domain and the middle domain (MD); M3: deletion of the LG domain, the hinge region, and α7–9 region in the MD domain; M4: deletion of the GTPase effector domain (GED); M5: E417A/K421A point mutations in the MD domain. h. Identification of GBP5 structural elements critical for inhibition of SARS-CoV-2 S protein glycosylation. Western blot analysis of 293T cells transfected with the indicated FLAG-GBP5 constructs and SARS-CoV-2 S protein for 48 h. Blots were probed with antibodies against FLAG and S protein. i. Identification of GBP5 structural elements critical for interaction with the OST complex. Western blot analysis of 293T cells transfected with the indicated FLAG-GBP5 constructs and MYC-tagged RPN1, RPN2, or DDOST for 48 h. Lysates were immunoprecipitated with anti-FLAG antibody and blots were probed with anti-MYC antibody. j. Schematic showing the proposed mechanism of GBP5 inhibition of SARS-CoV-2 S protein glycosylation. GBP5 interacts with the OST complex via accessory subunits and inhibits binding of S protein to the catalytic subunit STT3B.

GBP5 is localized in the cytoplasm whereas the major portions of RPN1, RPN2, and DDOST are in the ER lumen. To understand how GBP5 may interact with these ER luminal transmembrane proteins, we co-expressed FLAG-GBP5 with MYC-tagged RPN1, RPN2, and DDOST deleted for their C terminal cytoplasmic tails. GBP5 was immunoprecipitated with an anti-FLAG antibody, and we found that deletion of the cytoplasmic tails abolished the interaction between GBP5 and the OST subunits (Fig S3b). Collectively, these results indicate that GBP5 interacts with the OST accessory subunits RPN1, RPN2, and DDOST from the cytoplasmic side.

Interestingly, we did not detect direct binding between GBP5 and endogenous STT3A or STT3B (data not shown), indicating that GBP5 interacts directly with the three accessory subunits, but not the catalytic subunits, of the OST complex. Also of note, we did not observe co-immunoprecipitation of GBP5 with S protein (Fig S3c), suggesting that direct GBP5–S protein interactions are unlikely to be involved in the inhibition of S protein processing.

We next asked whether GBP5 might interfere with the interaction between the OST catalytic subunits and S protein substrate. We overexpressed FLAG-tagged SARS-CoV-2 S protein with GBP5 or GBP5-C583A in HEK 293T cells, and probed for the presence of endogenous STT3 subunits in anti-FLAG immunoprecipitates. Although the S protein– STT3B interaction was evident in cells overexpressing GBP5-C583A, the association was virtually abolished in cells expressing GBP5 (Fig 3f), suggesting that binding of GBP5 to the accessory subunits of OST hinders the ability of the OST catalytic subunits to interact with S protein.

Previous reports have suggested that the GTPase function of GBP5 is dispensable for inhibition of processing of other viral proteins^30^. To gain more insight into the structural and functional elements of GBP5 important for its interactions with the OST complex, we generated a series of GBP5 mutants and overexpressed them with SARS-CoV-2 S protein in 293T cells (Fig 3g). Consistent with existing evidence, deletion of the large GTPase catalytic (LG) domain had no effect on the ability of GBP5 to reduce S protein glycosylation or proteolysis (mutant M1, Fig 3g,h), and removal of the hinge (H) region connecting the LG domain to the middle domain (MD) similarly had no effect (M2, Fig 3G,H). The MD consists of five alpha helices, α7-α11, which together form two helical bundles bridged by α9 (Fig 3g,h)^46^. Removal of α7-α9 rescued S protein glycosylation, suggesting that this region contains residues or structural features essential for GBP5 interaction with the OST complex (M3, Fig 3g,h). Lastly, we found that truncating the C terminal GTPase effector domain (GED), while preserving the prenylation motif retained the WT GBP5 function, indicating that the GED domain is also dispensable for interfering with S protein glycosylation (M4, Fig 3g,h). Since the point mutations E417A and K421A, located at the C terminus of α9, have been shown to abolish the ability of GBP5 to inhibit HIV-1 particle infectivity^46^, we also examined their effects (M5, Fig 3g). Interestingly, this mutant did not inhibit S protein glycosylation or proteolysis (Fig 3h). Thus, the α7–α9 region, and E417 and K421 in particular, are essential for the function of GBP5 in S protein processing (Fig 3h).

To determine whether these mutations induced GBP5 loss-of-function by preventing GBP5 binding to OST, we overexpressed MYC-RPN2, -RPN1, or -DDOST and the FLAG-GBP5 WT or mutant constructs in HEK 293T cells and performed anti-FLAG immunoprecipitation followed by anti-MYC western blotting. The functional GBP5 mutants M1, M2, and M4 all associated with the three OST accessory subunits, as expected; however, we observed no associations between the nonfunctional mutant M3 or M5 GBP5s and RPN1, RPN2, or DDOST (Fig 3i and S3d). Collectively, these data support a model whereby GBP5 specifically interacts with the OST complex accessory subunits in a manner dependent on the GBP5 α7-9, thereby blocking S protein access to the OST catalytic site (Fig 3j). Taken together, our results suggest that GBP5 directly interacts with OST accessory subunits and blocks substrate access to the catalytic STT3 subunit.

### Scanning of SARS-CoV-2 S protein N-linked glycosylation sites revealed sequons essential for S protein maturation

The mechanism we delineated for GBP5 thus far, as well as our experiments using NGI-1, entails that proper N-linked glycosylation is intimately connected to viral glycoprotein maturation. To test whether S protein maturation indeed requires adequate N-linked glycosylation, we performed mutation scanning of twenty-two individual N-linked glycosylation sites in SARS-CoV-2 S and examined the effects on S protein proteolytic processing, trafficking, as well as virion assembly and release. We found that the majority of the 22 S protein N-to-Q mutations affected S protein processing, albeit to different extents, consistent with previous findings that select glycosylation sites contribute differentially to HIV envelope protein (env) maturation^47,48^. Eight mutations resulted in a significant reduction in proteolysis, and one mutation, N234Q, increased proteolysis (Fig 4a,b). Full-length S proteins bearing the eight mutations that reduced processing were completely sensitive to Endo H digestion (Fig 4c), suggesting that the reduced cleavage was primarily due to retention in pre-Golgi compartments. Most of the mutations that reduced S protein proteolysis by ≥50% yielded significantly lower titers of luciferase- or EGFP-encoding VSV pseudoviruses when tested with Calu-3 cells (Fig 4d,e), indicating a direct correlation between the reduction in S protein processing and assembly and secretion of infectious virions. Taken together, these data confirm the importance of N-linked glycosylation for S protein maturation, and shows that it is possible for host factors that interfere with S protein glycosylation to suppress the generation of infectious virions.

**Figure 4.**
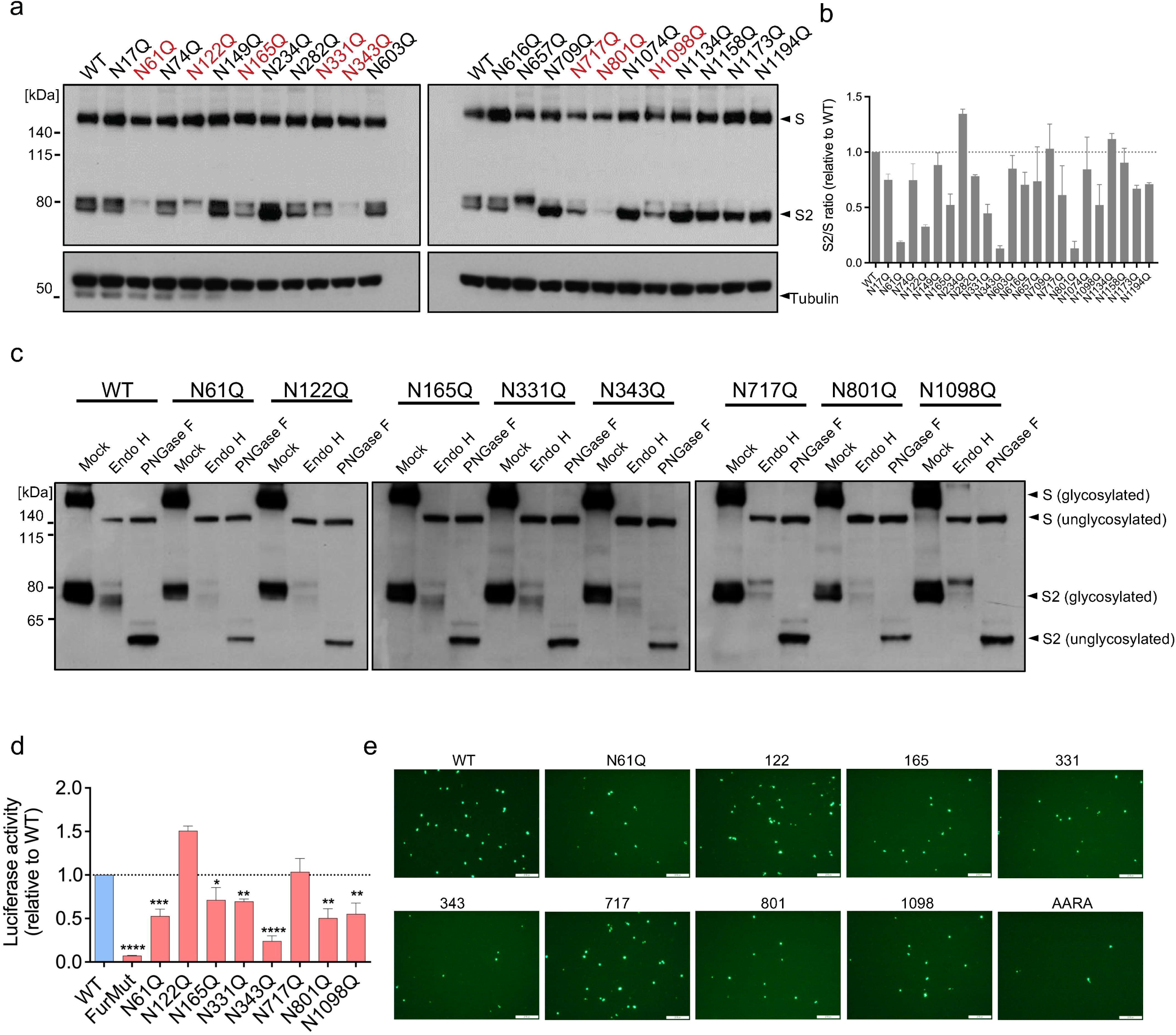
Scanning mutagenesis identifies SARS-CoV-2 S protein N-linked glycosylation sites important for infection. a, b. (A) Western blot analysis of 293T cells transfected for 48 h with the indicated S protein glycosylation site mutants. Blots were probed with antibodies against S protein or tubulin (loading control). Mutants showing the strongest cleavage inhibition are shown in red. (B) Ratio of S2 subunit to full-length S protein expressed by the cells shown in (A). Mean ± SD of two independent experiments. c. Western blot analysis of 293T cells transfected for 48 h with the indicated S protein mutant constructs. Cell lysates were digested with Endo H and PNGase F for 1 h. Blots were probed with anti-S2 antibody. d, e. Mutation of S protein glycosylation sites impairs infectivity of VSV pseudoviruses. Luciferase- or EGFP-encoding (I) VSV pseudoviruses expressing the indicated WT or mutated SARS-CoV-2 S proteins were produced in 293T cells. After 24 h, the supernatants were collected and used to infect Calu-3 cells for 2 h. Infectivity of VSV pseudoviruses was assessed by luciferase assays (D) or fluorescence microscopy (E). Mean ± SD of n = 3. *p < 0.05, **p < 0.01, ***p < 0.001, ****p < 0.0001 by Student’s t-test. Scale bar, 100 μm.

### GBP5 inhibits protein glycosylation and trafficking in multiple viruses

GBP5 has been shown to inhibit the maturation of other viral glycoproteins, including influenza A virus (IAV) HA and HIV gp160, both of which are cleaved by host proprotein convertases in the Golgi apparatus^29^. To determine whether these GBP5 effects are also mediated via interference with protein glycosylation and ER-to-Golgi trafficking, we analyzed HEK 293T cells co-transfected with GBP5 or GBP5-C583A and FLAG-tagged IAV HA or HIV gp160. Indeed, overexpression of GBP5 inhibited the proteolytic processing and decreased MWa of both full-length HA (HA_0_) and full-length gp160 compared with control cells, whereas GBP5-C583A overexpression did not (Fig 5a,b). GBP5 overexpression also decreased the proteolytic processing of MERS-CoV S protein (Fig S4a), as well as the MWa of SARS-CoV S protein (Fig S4b), which does not undergo proteolytic processing during maturation. These data suggest that the effects of host GBP5 on the processing and maturation of viral glycoproteins are not restricted to SARS-CoV-2 and may be of importance to a wide variety of virus species and families.

**Figure 5.**
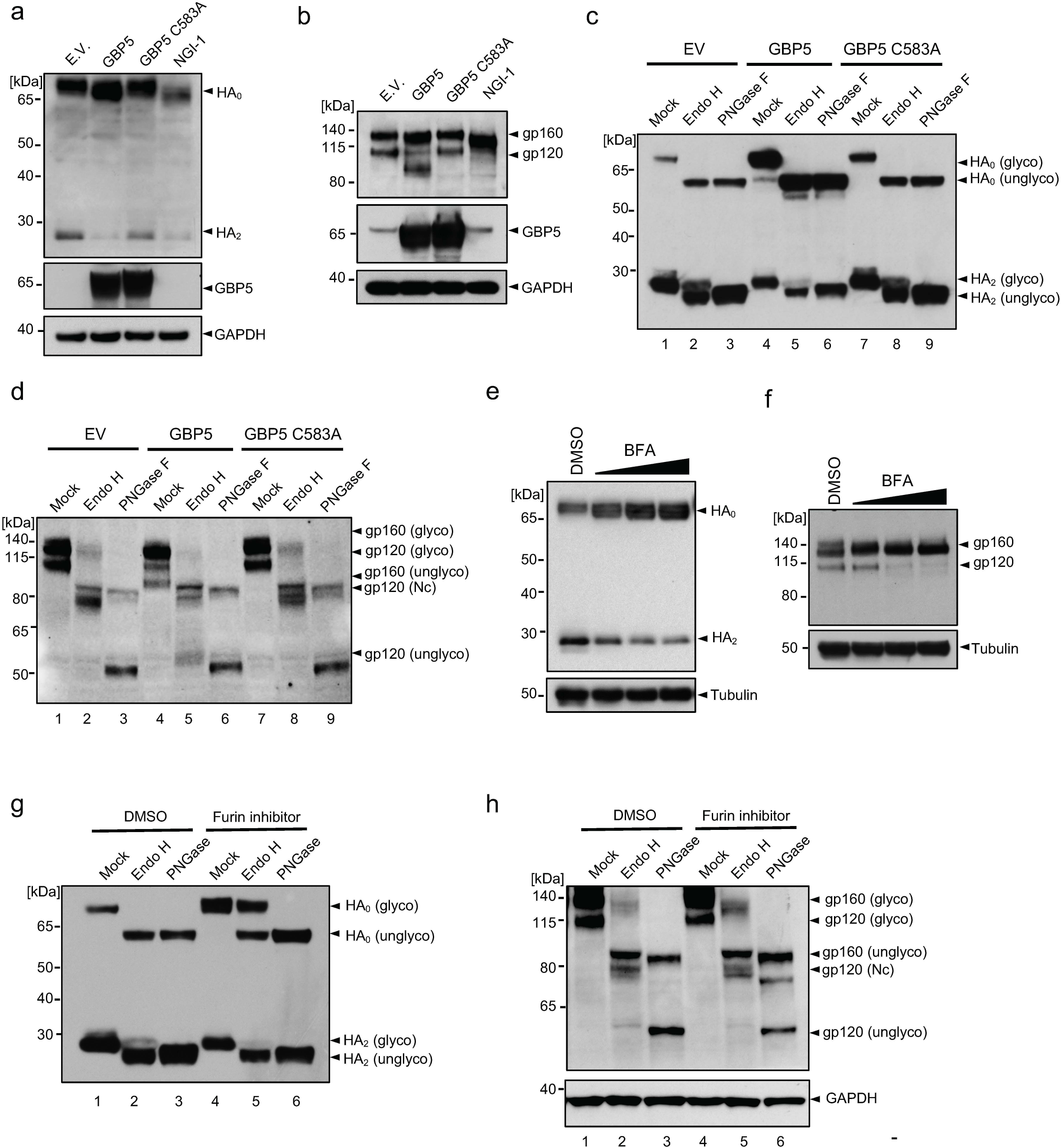
GBP5 inhibits glycoprotein processing in multiple viruses. a, b. GBP5 inhibits IAV HA and HIV gp160 glycosylation and proteolytic processing. Western blot analysis of 293T cells transfected with empty vector (EV), GBP5, or GBP5-C583A and co-transfected with FLAG-tagged IAV HA for 24 h (A) or with HIV pNL4-3 (encoding HIV NL4-3 provirus gp160) for 48 h (B). Blots were probed with antibodies against GBP5, GAPDH (loading control) and either FLAG (A) or gp120 (B). Transfected cells treated with NGI-1 for 24 h were included as positive controls. c, d.. Sensitivity of IAV HA and HIV gp160 to Endo H and PNGase F digestion. Western blot analysis of cell lysates prepared as in (A, B) after digestion for 1 h with Endo H or PNGase F. Blots were probed with antibodies against FLAG (C) or gp120 (D). e, f. Blocking ER-to-Golgi trafficking inhibits proteolytic processing of IAV HA and HIV gp160. Western blot analysis of 293T cells transfected with FLAG-IAV HA for 18 h (E) or with HIV pNL4-3 for 42 h (F) and then treated with DMSO or 0.012, 0.05, or 0.2 μM BFA for 6 h. Blots were probed with antibodies against FLAG (E), gp120 (F), and tubulin (loading control). g, h. Inhibition of furin-mediated SARS-CoV-2 S protein cleavage does not resemble GBP5 overexpression. Western blot analysis of lysates of 293T cells transfected with FLAG-IAV HA (G) or HIV pNL4-3 (H) and incubated with DMSO or 100 μM CMK for 24 h (HA) or 48 h (pNL4-3). Cell lysates were digested with Endo H or PNGase F for 1 h. Blots were probed with antibodies against FLAG (G), gp120 (H), and GAPDH (loading control).

To determine whether the observed GBP5-dependent reductions in MWa of IAV HA and HIV gp160 were indeed due to a reduced quantity of N-linked glycans, we digested lysates of transfected HEK 293T cells with Endo H or PNGase F. Indeed, PNGase F treatment generated fragments with electrophoretic mobility corresponding to unglycosylated HA_0_ and HA_2_ (cleaved HA) under GBP5 overexpression (Fig 5c, lanes 3, 6, 9), showing that GBP5 interferes with HA_0_ glycosylation. Similar to SARS-CoV-2 S protein, HA_0_ was completely sensitive to Endo H (Fig 5c, lanes 2, 5, 8), suggesting that HA_0_ harbored high mannose-type N-linked glycans and localized in the ER. HA_2_ was mostly sensitive to Endo H but still possessed partial resistance (Fig 5c, lanes 2, 5, 8), suggesting that cleaved HA had gone through complex glycan modification in the Golgi apparatus. Compared to control and GBP5-C583A overexpression, overexpression of wild-type GBP5 led to an accumulation of HA_0_, which localizes to the ER, and a depletion of HA_2_, which trafficked past Golgi compartments (Fig 5c, lanes 2, 5, 8). Thus, overexpression of GBP5 dampened ER-to-Golgi trafficking of HA. Because Endo H also removed the decrease in the MWa of HA_0_ under GBP5 overexpression (Fig 5c, lanes 2, 5, 8), disruption of HA N-linked glycosylation should have taken place where N-linked glycans on HA were still high mannose-type. These observations suggest that GBP5 decreases HA N-linked glycosylation and trafficking to post-Golgi compartments.

In the case of HIV env, both full-length env (gp160) and cleaved env (gp120) demonstrated the same MWa after PNGase F digestion (Fig 5d, lanes 3, 6, 9), suggesting GBP5 reduced the amount of N-linked glycans on gp160 and the majority of gp120. gp160 expressed with an empty vector or GBP5-C583A was largely Endo H-sensitive but showed residual Endo H-resistant bands (Fig 5d, lane 2, 8). This suggests that some gp160 molecules had been trafficked to the Golgi compartments but had not been cleaved by furin. The residual Endo H resistant band of gp160 vanished under GBP5 overexpression compared to control and GBP5-C583A overexpression (Fig 5d, lane 5), suggesting that overexpression of GBP5 inhibited ER-to-Golgi trafficking of gp160.

In line with expectation, blocking ER exit by BFA abolished proteolytic processing of HA and gp160 (Fig 5e, f), showing that secretory trafficking is required for these glycoproteins to properly mature. Given previous reports proposing that GBP5 interferes with HA and gp160 processing through inhibiting furin activity, we used CMK to probe the potential difference between GBP5 overexpression and furin inhibition. Consistent with our model worked out using S protein, CMK treatment decreased HA and gp160 cleavage without lowering the MWa of either (Fig S4c, d). The accumulation of HA_0_ due to CMK was accompanied by a strong Endo H-resistant HA_0_ signal (Fig 5g, lane 5), whereas HA_0_ in GBP5-overexpressing cells was completely sensitive to Endo H. In the case of HIV gp160, CMK treatment increased the band intensity of Endo H-resistant gp160 (Fig 5h), contrary to GBP5 overexpression which sensitized gp160 to Endo H. These data show that GBP5 restricts HA and gp160 maturation primarily by interfering with their N-linked glycosylation, leading to retention in pre-Golgi compartments, not by inhibiting furin activity.

### Pharmacological inhibition of OST inhibits maturation of multiple viruses

Having shown that inhibition of OST with NGI-1 largely mimicked GBP5 by dose-dependently reducing the glycosylation and cleavage of SARS-CoV-2 S protein and production of SARS-CoV-2 pseudovirus (Fig 2f, 6a, and S5a)^36^, we further characterized the antiviral potential of NGI-1. First, we examined the impact of incubating Calu-3 and Caco-2 cells with NGI-1 after infection with authentic SARS-CoV-2 WT. Quantification of viral RNA in both the infected cells and the culture supernatants showed that NGI-1 dose-dependently inhibited SARS-CoV-2 WT infection of Calu-3 and Caco-2 cells at 50% inhibitory concentrations (IC_50_) of 3.24 μM and 3.39 μM, respectively (Fig 6b-e, S5b, c). Importantly, we also found that NGI-1 could inhibit infection of human stem cell-derived human lung organoids, a more physiologically relevant model^49^, with SARS-CoV-2 WT (Fig 6f,g), and of Calu-3 cells with a clinical isolate of SARS-CoV-2 B.1.1.529 (Omicron) (Fig 6h,i). Finally, we examined 293T cells transfected with VSV pseudoviruses bearing S proteins from B.1.351 (Beta), B.1.617.2 (Delta), and B.1.1.529 (Omicron), and found that incubation with NGI-1 dose-dependently inhibited glycosylation and proteolysis of S proteins from each strain as well as production of infectious pseudoviruses (Fig 6j,k, S5d– g).

**Figure 6.**
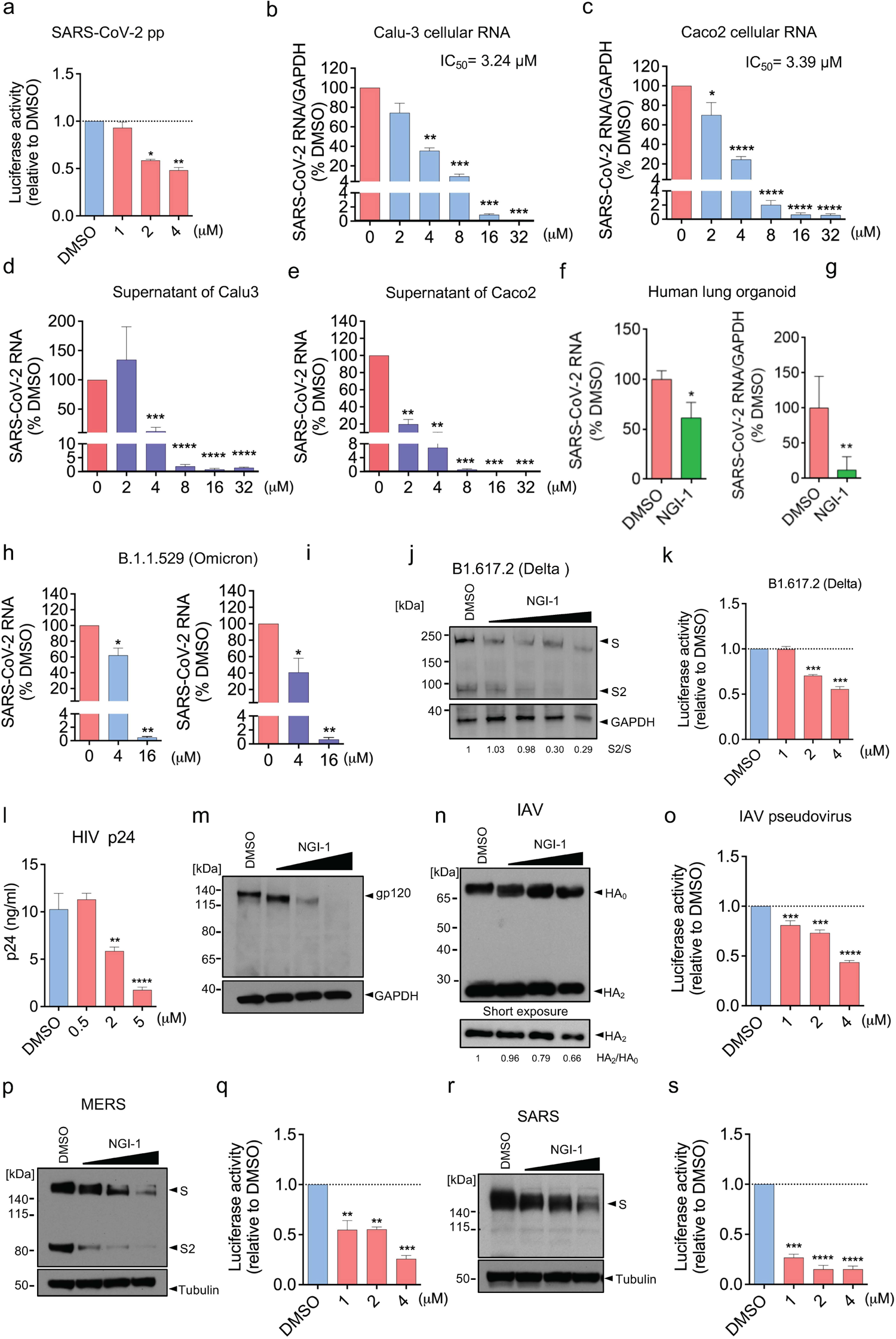
Pharmacological inhibition of OST suppresses maturation of multiple viruses. a. Inhibition of OST decreases SARS-CoV-2 pseudovirus production. Luciferase-expressing SARS-CoV-2 S protein VSV pseudovirus was produced in 293T cells treated with DMSO or NGI-1. After 24 h, pseudovirus-containing supernatants were collected and used to infect Calu-3 cells for 2 h. Infectivity was measured by luciferase assay. Mean ± SD of n = 3. *p < 0.05, **p < 0.01 by Student’s t-test. b-e. Calu-3 (B, D) and Caco-2 (C, E) cells were infected with SARS-CoV-2 USA-WA1/2020 at a MOI of 0.1 and treated with DMSO or NGI-1 for 24 h (Calu-3) or 48 h (Caco-2). Viral RNA in the cells (B, C) and supernatants (D, E) was quantified by RT-qPCR. Mean ± SD of n = 3. *p < 0.05, **p < 0.01, ***p < 0.001, ****p < 0.0001 by Student’s t-test. f, g. Human lung organoids (60 days) were pretreated with DMSO or 5 μM NGI-1 for 2 h and then infected with SARS-CoV-2 USA-WA1/2020 at an MOI of 0.1 for 48 h. Viral RNA in the supernatants (F) or organoids (G) was quantified by RT-qPCR. Mean ± SD of n = 2, representative of 3 biological replicates. *p < 0.05, **p < 0.01 by Student’s t-test. h, i. Calu-3 cells were infected with SARS-CoV-2 B.1.1.529 (Omicron) at an MOI of 0.1 and then treated with DMSO or NGI-1 for 48 h. Viral RNA in cells (H) and supernatants (I) was quantified by RT-qPCR. Mean ± SD of n = 3. **p < 0.01, ***p < 0.001 by Student’s t-test. j, k. Luciferase- and SARS-CoV-2 B.1.617.2 (Delta) S protein-expressing VSV pseudovirus was produced in 293T cells treated with DMSO or 1, 2, or 4 μM NGI-1. After 24 h, cells and supernatants were collected. (J) Cell lysates were subjected to western blot analysis with antibodies against S protein or GAPDH (loading control). (K) Supernatants were used to infect Calu-3 cells for 2 h and infectivity was quantified by luciferase assay. Mean ± SD of n = 3. **p < 0.01, ***p < 0.001 by Student’s t-test. l, m.. MT4 cells were infected with HIV-1 LAI at an MOI of 0.01 and treated with DMSO or NGI-1 for 48 h. (L) Cell supernatants were analyzed for p24 levels by ELISA. (M) Cell lysates were analyzed by western blotting with antibodies against gp120 and GAPDH (loading control). Mean ± SD of n = 4. **p < 0.01, ****p < 0.0001 by Student’s t-test. n, o. Luciferase- and IAV HA-expressing VSV pseudovirus was produced in 293T cells treated with DMSO or 1, 2, or 4 μM NGI-1. After 24 h, cells and supernatants were collected. (n) Cell lysates were subjected to western blot analysis with an antibody against HA. (o) Supernatants were used to infect A549 cells for 2 h and infectivity was measured by luciferase assay. Mean ± SD of n = 3.***p < 0.001, ****p < 0.0001 by Student’s t-test. p-s. Luciferase-expressing VSV pseudoviruses were produced in 293T cells transfected with MERS-CoV (p, q) or SARS-CoV (r, s) S proteins and treated with DMSO or 1, 2, or 4 μM NGI-1. After 24 h, cells and supernatants were collected. (p, r) Cell lysates were subjected to western blot analysis with antibodies against S protein or tubulin (loading control). (q, s) Supernatants were used to infect Calu-3 cells for 2 h, and infectivity was quantified by measuring luciferase activity. Mean ± SD of n = 3. **p < 0.01, ***p < 0.001, ****p < 0.0001 by Student’s t-test.

Because GBP5 overexpression inhibited the maturation of SARS-CoV-2, MERS-CoV, and SARS-CoV S proteins and additionally of IAV HA and HIV gp160, we considered that NGI-1 might exhibit broad-spectrum antiviral activity. To test this, we infected the human CD4+ T cell line MT4 with HIV-1 LAI and then treated the cells with NGI-1. HIV p24 and viral RNA levels in the culture supernatants, as well as cellular viral RNA levels, were all reduced by NGI-1 treatment (Fig 6l, S5h-j), confirming that NGI-1 could suppress HIV replication. Moreover, NGI-1 also reduced gp160/gp120 glycosylation and processing (Fig 5b and 6m), similar to its effects on SARS-CoV-2 S protein processing. To confirm that NGI-1 inhibits HIV replication at the later step of virion maturation, 293T cells were treated with NGI-1 during the packaging of luciferase-expressing HIV LAI pseudoviral particles. Infectivity was tested by application of the pseudovirus-containing 293T cell supernatants to MT4 cells followed by measurement of luciferase activity. Indeed, NGI-1 treatment during pseudovirus packaging significantly inhibited the infectivity of MT4 cells (Fig S5k), demonstrating that NGI-1 inhibits HIV virion assembly and maturation. Finally, we produced VSV pseudoviruses bearing IAV HA, MERS-CoV S protein, and SARS-CoV S protein in 293T cells treated with NGI-1 and then analyzed HA and S protein processing, as well as infectivity of pseudovirus supernatants on A549 (IAV HA) or Calu-3 cells (S proteins). We found that NGI-1 inhibited HA glycosylation and cleavage and also decreased the infectivity of HA-pseudotyped VSV (Fig 6n,o). In addition, similar effects of NGI-1 treatment were observed for processing of MERS-CoV S protein (Fig 6p) and infectivity of VSV pseudovirus (Fig 6q), as well as decreased SARS-CoV S protein MWa (Fig 6r) and VSV pseudovirus infectivity (Fig 6s). Taken together, these results show that inhibition of OST complex activity by NGI-1 inhibits viral protein maturation and virion infectivity for a range of viruses.

## Discussion

Viral protein maturation and virion assembly are dependent on host protein glycosylation pathways. In this study, we described a novel mechanism of antiviral activity for the ISG GBP5 via interference with OST-mediated transfer of N-glycan precursors to viral proteins, thereby promoting their ER retention and inhibiting further processing. We showed that GBP5 directly interacts with OST accessory subunits and blocks binding of the OST catalytic subunits to SARS-CoV-2 S protein. The consequences of this disruption of N-linked glycosylation included S protein misfolding, aggregation, and retention in the ER, and subsequent impairment of infectious virion assembly and secretion (Fig 7). Moreover, we showed that pharmacological inhibition of the OST complex exhibited similar effects of GBP5 overexpression on SARS-CoV-2, SARS-CoV, MERS-CoV, IAV, and HIV protein glycosylation and processing and virion assembly and release, further confirm that OST-specific inhibitors might exhibit broad antiviral activity. In previous studies, GBP5 has been shown to restrict release of lentivirus pseudotyped with surface glycoproteins of rabies virus, European bat lyssavirus, Marburg virus, measles virus, and murine leukemia viruse^29^. Therefore, mimicking the antiviral mechanism of GBP5 might represent a promising broad antiviral strategy.

**Figure 7.**
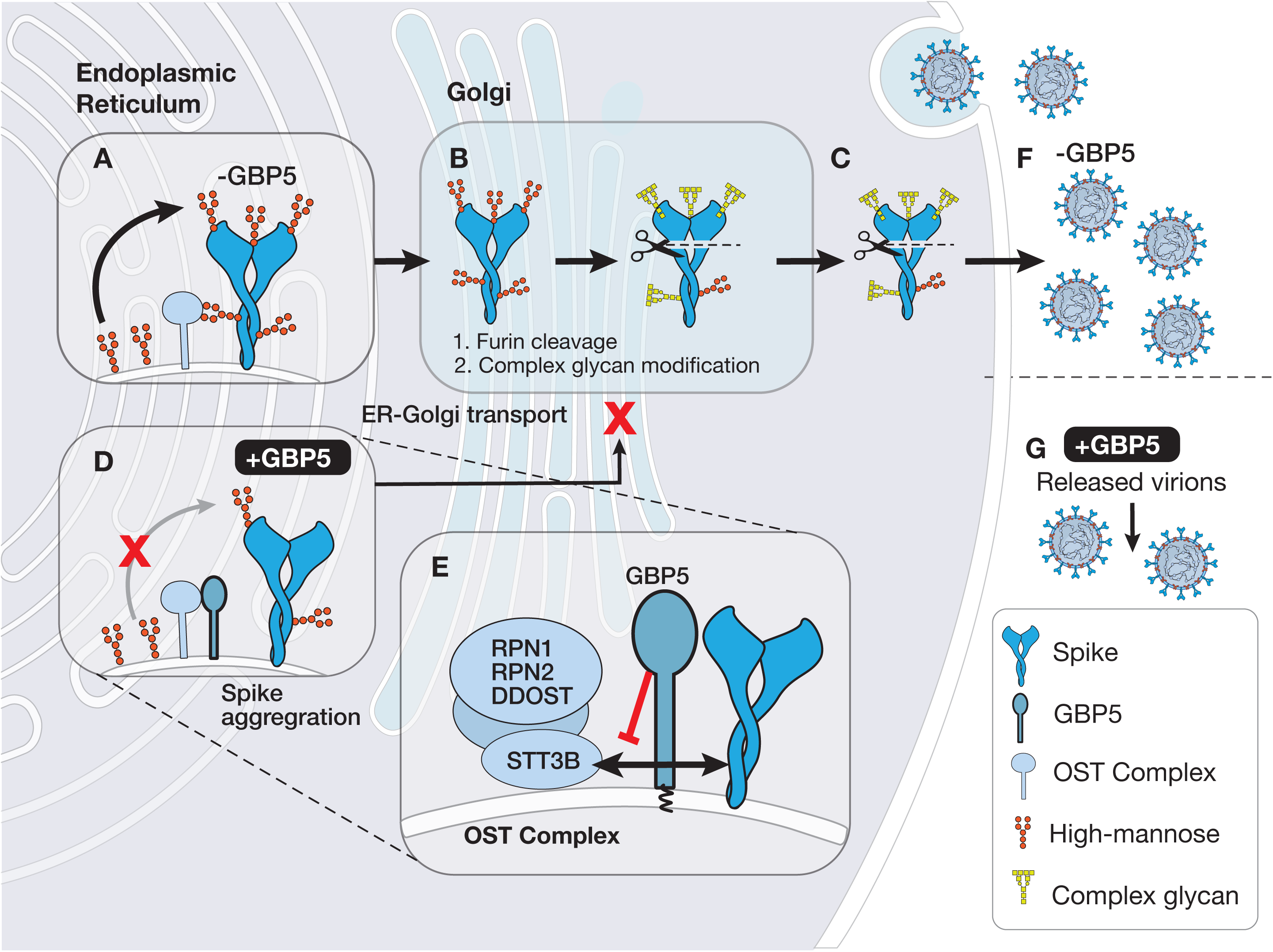
Proposed model for GBP5 involvement in the maturation of viral S proteins. Viral S protein is glycosylated with high mannose-type glycans by the OST complex in the ER (A) and transported to the Golgi apparatus for complex glycan modification and furin cleavage (B), resulting in export of properly glycosylated and cleaved S protein (C), which is assembled into new virions and released from the cell (F). In the presence of high levels of GBP5, the interaction between GBP5 and the OST complex prevents proper S protein glycosylation, leading to misfolding and aggregation of S protein in the ER (D). As a result, transport of S protein to the Golgi apparatus is reduced and furin cleavage is blocked (B), thus inhibiting the assembly and release of virions (G). Mechanistically, GBP5 inhibits OST by blocking its interaction with the accessory subunits RPN1, DDOST and RPN2, thereby preventing the interaction between S protein and the OST catalytic subunit STT3B (E).

GBP5 has previously been demonstrated to have several different mechanisms and exhibit antiviral activity against multiple viruses including HIV, influenza A, rabies, as well as in European bat lyssa. During RSV infection, GBP5 was shown to decrease viral small hydrophobic (SH) protein levels, thereby inhibiting RSV replication^50^. GBP5 is also induced during HIV infection of human macrophage and CD4^+^ T cells and was initially identified and validated as an HIV restriction factor through a genome-wide evolutionary profiling study^30^. GBP5 has previously been proposed to act by inhibiting gp160 cleavage via binding to and inhibition of furin^29^, which contrasts with the results of the present study. However, there are several possible weaknesses to this model. First, GBP5 was shown to inhibit the infectivity of HIV pseudotyped with Lassa virus glycoprotein (LASV GP)^29^. However, LASV GP is cleaved by a non-basic proprotein convertase, SKI-1/S1P, which has an amino acid sequence and structure quite different from those of furin. Thus, it remains unclear whether GBP5 and S1P can actually interact^51^. Second, direct inhibition of furin by GBP5 would not explain the change in MWa of gp160. Furthermore, although evidence that GBP5 specifically affects furin activity using purified furin or in cells overexpressing furin has been reported, such evidence has not been observed for endogenously expressed furin in cells, and we found that GBP5 overexpression did not alter the endogenous furin activity like furin inhibitor CMV-RVKR^29,52^.Thus, the results of the present study allow us to propose a model in which GBP5 affects the maturation of any viral protein that is glycosylated by OST in the ER and undergoes maturation by Golgi-resident proprotein convertases, which simultaneously explains the reduction in MWa and proteolytic processing of OST substrates observed in GBP5-overexpressing cells. Our results therefore suggest that GBP5 exerts antiviral activity primarily by inducing defects in viral protein N-linked glycosylation and trafficking.

We added to the accumulating evidence of the feasibility of therapeutic targeting of the OST complex by using the small molecule inhibitor NGI-1, which potently suppressed SARS-CoV-2 WT infection of human lung and colon adenocarcinoma cells lines with IC_50_ values in the low micromolar range. Mechanistically, NGI-1 inhibits OST by directly binding to the catalytic STT3 subunits^32^, which slightly differs from the mechanism of action of GBP5 identified here. This may explain the greater magnitude of reduction in protein MWa in NGI-1-treated versus GBP5-overexpressing cells observed in our study, and the ability of NGI-1 to induce ER stress might account for the dose-dependent decrease in S protein levels observed here^36^. The roles of the OST accessory subunits in glycan transfer are poorly understood, but loss of expression of specific accessory subunits has been associated with loss of N-linked glycans on distinct substrates^53–55^. Thus, by binding solely to the OST accessory subunits, GBP5 may modulate the glycosylation of a more restricted panel of substrates and result in a more subtle phenotype compared with NGI-1. Our work suggests that modulators of OST could not only have potential therapeutic utility but also provide useful tools for glycosylation research. Additional structural information about the GBP5–OST interaction would serve as the basis for understanding the structure–activity relationship to inform the design and development of small molecule mimetics.

In conclusion, the results of this study contribute to our understanding of the interplay between the innate antiviral immune response and the function of ISGs during viral maturation. We uncovered a novel strategy by which host cells may combat viral infection, thereby shedding light on the importance of enzymes and cofactors involved in viral protein glycosylation during the late stages of viral infection. Further studies are also likely to reveal new potential strategies for the development of novel broad-spectrum antiviral drugs.

## Materials and Methods

### Cell culture, viruses, and reagents

All studies were performed based on the approved IRB protocols by the University of California (UCSD), San Diego. The human carcinoma cell lines Calu-3 and Caco-2 were purchased from ATCC and were cultured in MEM medium supplemented with 1×NEAA, 100 IU/mL penicillin–streptomycin, and 20% FBS. HEK293T (293T), A549 and Vero cells were maintained in our lab and were cultured in DMEM medium supplemented with 10% FBS. MT4 cell was cultured in RPMI medium 1640 with 10% FBS. 293T cells transduced with lentivirus encoding human ACE2 and TMPRSS2 was maintained in DMEM medium with 10% FBS and 2 ug/ml puromycin. Brefeldin A was purchased from BioLegend; kifunensine, swainsonine, decanoyl-RVKR-CMK, and NGI-1 were purchased from Cayman Chemical; Endo H and PNGase F were purchased from New England Biolabs; The DNA fragments encoding spike of SARS-CoV-2 strains USA-WA1/2020 (WT), B. 1. 351 (Beta) were inserted into pLVX-puro and C-terminal was tagged with Flag, no organelle localization signal was included in this vector. The spike plasmids of B.1.617.2 (Delta), and B.1.1.529 (Omicron) were obtained from InvivoGen. The plasmids of SARS-CoV (CUHK-W1) spike, MERS-CoV (HCoV-EMC/2012) spike, IAV H5N1 (Thailand/KAN-1/2004) HA and NA were purchased from Sino Biological. All the plasmids of viral glycoproteins above were codon-optimized.

### Pseudovirus production and infection assays

EGFP- or luciferase-expressing VSV pseudoviruses were generated as previously described^56^. Briefly, BHK-21/WI-2 cells were infected with vTF7-3 for 45 min and then transfected with pVSV-EGFP-dG (Addgene 31842) or pVSV-FLuc-dG and pBS vectors encoding the N, P, L, and G proteins of VSV (Kerafast) at a mass ratio of 3:3:5:1:8 (pVSV-EGFP-dG/pVSV-Fluc-dG: pBS-N: pBS-P: pBS-L: pBS-G). Transfection was performed with Lipofectamine 3000 (Invitrogen) according to the manufacturer’s instructions. At 48 h after transfection, BHK-21/WI-2 culture supernatants were filtered (0.22 μm) and used to inoculate 293T cells that had been transfected 24 h previously with pMD2.G (Addgene 12259). At 24 h post-inoculation, 293T culture supernatants were collected and stored at −80°C.

For the production of SARS-CoV-2, SARS-CoV, and MERS-CoV pseudoparticles, 293T cells were transfected with plasmids expressing the S proteins of SARS-CoV-2 USA-WA1/2020 (WT), Beta variant, Delta variant, Omicron variant, SARS-CoV CUHK-W1, and MERS-CoV (HCoV-EMC/2012) for 24 h and infected with VSV pseudovirus for 1 h. Nineteen amino acids at the C terminus of SARS-CoV-2 and SARS-CoV S proteins were deleted for better incorporation into the viral envelope^57^. The infected 293T cells were washed four times with medium and incubated for an additional 24 h in the presence of NGI-1 or DMSO. The culture supernatants were collected, centrifuged to remove cell debris, and stored at −80°C. For the infectivity assays, Calu-3 cells were incubated with EGFP-expressing or Luc-expressing pseudoviruses for 2 h and then analyzed by fluorescence microscopy or luciferase assays, respectively.

### Virus-like particle (VLP) production

SARS-CoV-2 VLP were generated as previously described^31,32^. In brief, plasmids encoding the viral structural proteins S, M, E, and N were transfected into 293T cells for 48 h transfection. The supernatant was collected, passed through a 0.45 μm filter, and concentrated over 20% sucrose by ultracentrifugation at 28,000 rpm for 3 h at 4°C. For analysis, the pellets were resuspended in 50 μL 2x SDS buffer and subjected to SDS-PAGE.

### SARS-CoV-2 infection

293T/ACE2/TMPRSS2 cells were transfected with empty vector (EV), GBP5, or GBP5-C583A for 24 h and then infected with SARS-CoV-2 USA-WA1/2020 at the indicated MOIs for 1 h at 37°C. The infected cells were washed and incubated in complete medium for 24 h before analysis. Calu-3 and Caco-2 cells were infected with SARS-CoV-2 USA-WA1/2020 at a MOI of 0.1 for 1 h at 37°C. The cells were washed and incubated in fresh medium with DMSO or NGI-1 for 24 h (Calu-3) or 48 h (Caco-2). Calu-3 cells were infected with SARS-CoV-2 Omicron at a MOI of 0.1 and incubated with DMSO or NGI-1 for 48 h. SARS-CoV-2 WA1 (USA-WA1/2020) was acquired from BEI (cat # NR-52281) and propagated on TMPRSS2-VeroE6 cells. SARS-CoV-2 BA.1/Omicron (hCoV-19/USA/CA-SEARCH-59467/2021) was isolated on TMPRSS2-VeroE6 cells from a patient nasopharyngeal swab sample under UC San Diego IRB #160524 with sequence deposited at GISAID (EPI_ISL_8186377). Viral stocks were verified by deep sequencing.

### Culture and maintenance of human-induced pluripotent stem cell (hiPSC)-derived 3D lung organoids

All studies related to human tissues and samples were performed through approved IRB protocols by the University of California (UCSD), San Diego. Human-induced pluripotent stem cells (hiPS11-10, Cell Application) were cultured on basement membrane growth factor-reduced Matrigel-coated plates in mTeSRTM1 medium at 37°C. hiPSCs were differentiated into human lung organoids and characterized as previously established by our group and others^49,58^. In brief, ∼50–75% confluent undifferentiated hiPSC colonies were differentiated stepwise with specific medium into definitive endoderm (RPMI containing 0.2–2% FBS and 100 ng/mL Activin A), anterior foregut endoderm (Advanced DMEM/F12 medium containing N2-1x, B27-1x, 10 mM HEPES buffer, 2 mM L-glutamine, penicillin–streptomycin-1x, 0.4 μM monothioglycerol, 0.25% BSA, 50 μg/mL ascorbic acid, 10 μM SB431542, 200 ng/mL Noggin, 1 μM SAG, 500 ng/mL FGF4, and 2 μM CHIR-99021), and lung organoids (DMEM/F12 medium containing N2-1x, B27-1x, 10 mM HEPES buffer, 2 mM L-glutamine, penicillin–streptomycin-1x, 0.4 μM monothioglycerol, 0.25% BSA and 50 μg/mL ascorbic acid, 10 ng/mL each of FGF7, FGF10, EGF, and VEGF/PIGF, 3 μM CHIR, 50 nM ATRA, 100 μM cAMP, 100 μM IBMX, and 50 nM dexamethasone). Lung organoids were maintained by re-embedding in Matrigel every 2– 3 weeks. Lung organoids were examined for expression of ACE2, TMPRSS2, neuropillin, and the alveolar cell epithelial markers SFTPC, SFTPB, and HOPX by qPCR^49^ and were employed at 60 days for experiments.

### Lung organoid infection

Infectious units of SARS-CoV-2 USA-WA1/2020 were quantified using a plaque assay with Vero E6 cells. Lung organoids were pretreated with NGI (5 µM) for 2 h and then infected with SARS-CoV-2 at a MOI of 0.1 for 2 h at 37°C as described previously^44^. The organoids were washed and fresh medium containing 5 μM NGI-1 was added. After 48 h, the supernatants and organoids were collected.

### Assessment of glycoprotein sensitivity to endoglycosidases

Cell lysates were digested for 1 h with Endo H (P0702, NEB) or PNGase F (P0704, NEB) according to the manufacturer’s instructions. Upon completion of digestion, proteins were precipitated by the addition of four volumes of acetone (prechilled at −20°C) followed by incubation for 30 min at −20°C. The samples were centrifuged at 15,000 g at 4°C for 15 min, the supernatants were discarded, and the precipitated proteins were dried at room temperature and analyzed by SDS-PAGE.

### Lentiviral production and establishment of stable cell lines

To generate lentiviral particles, 293T cells were seeded in 6-well plates and co-transfected with pMD.2G (0.6 μg), psPAX.2 (1.2 μg), and pLVX encoding ACE2 or TMPRSS2 (1.8 μg) using Lipofectamine 3000, according to the manufacturer’s instructions. At 48 h post-transfection, the culture supernatants were collected and added to 293T cells in the presence of 10 μg/mL polybrene. Cells were centrifuged at 750 g for 45 min to enhance transduction efficiency. At 24 h post-transduction, the medium was replaced with fresh medium supplemented with 2 μg/mL puromycin and the cells were maintained in 2 μg/mL puromycin-containing medium to select for transduced cells.

### HIV infection of MT4 cells

MT4 cells were resuspended in DMSO- or NGI-1-containing medium and infected with HIV-1 LAI strain at a MOI of 0.01. At 24 h post-infection, the medium was exchanged for fresh medium containing DMSO or NGI-1 and the cells were incubated for an additional 72 h. The culture supernatants were collected and p24 levels were quantified by ELISA or viral RNA levels were quantified by one-step RT-qPCR. The cells were also collected and subjected to western blot analysis of gp160 or RT-qPCR analysis of viral RNA. For HIV pseudovirus infection, 293T cells were transfected with pNL4-3.Luc.R-E, which encodes full-length HIV-1 proviral DNA without envelope, and an envelope-expressing plasmid in the presence of NGI-1 or DMSO. At 48 h post-infection, the supernatant containing HIV pseudovirus was collected and added to MT4 cells. After 48 h, the cells were collected and infectivity was determined by luciferase assay according to the manufacturer’s instructions (E1500, Promega).

### Cell viability assay

Cell viability assays were performed with CellTiter-Glo 2.0 (Promega) according to the manufacturer’s instructions.

### RNA extraction, qPCR, and one-step RT-qPCR

RNA was extracted from cells and culture supernatants with TRIzol and TRIzol LS, respectively and purified with Direct-zol RNA Purification Kits (Zymo). Viral RNA was quantified by qPCR using primers targeting the nucleocapsid: SARS-CoV-2-N (Forward: CACATTGGCACCCGCAATC; Reverse: GAGGAACGAGAAGAGGCTTG). GAPDH was analyzed as an internal control mRNA (Forward: GGCCTCCAAGGAGTA-AGACC; Reverse: AGGGGTCTACATGGCAACTG).

### Native PAGE

Native PAGE was performed using the Novex NativePAGE Bis-Tris Gel System (Thermo Scientific) according to the manufacturer’s protocol. In brief, 293T cells co-transfected with SARS-CoV-2 S protein and EV, GBP5, or GBP5-C58A were lysed in sample buffer containing 2% digitonin and 0.5% G250 sample additive. Samples were centrifuged at 13,300 rpm at 4°C for 30 min, and equal amounts of protein were separated on a NativePAGE 3–12% Bis-Tris gel. Proteins were transferred to a PVDF membrane and S protein oligomerization was detected by western blot analysis.

### Immunoprecipitation

Cells were lysed with Pierce IP Lysis Buffer (Thermo Scientific) and centrifuged. Supernatants (2 mg protein) were removed and incubated overnight at 4°C with Dynabeads Protein G (Thermo Scientific) coated with anti-FLAG antibody or IgG heavy chain according to the manufacturer’s instructions. The next day, the beads were collected by centrifugation and washed using one of two methods. For co-immunoprecipitation of FLAG-GBP5 or FLAG-GBP5-C583A with MYC-tagged ectopic or endogenous RPN2, RPN2, and DDOST, beads were washed once with Pierce IP Lysis Buffer, twice with Pierce IP Lysis Buffer supplemented with 500 mM NaCl, and once with a low salt buffer. For co-immunoprecipitation of FLAG-SARS-CoV-2 S protein with endogenous STT3B or STT3A, beads were washed three times with Pierce IP Lysis Buffer. Immunocomplexes and aliquots of input lysates were subjected to SDS-PAGE followed by western blot analysis.

### Construction of SARS-CoV-2 S protein glycosylation mutants

Codon-optimized full-length SARS-CoV-2 S protein gene was cloned into Topo cloning vector and 22 N-glycosylation sites were mutated individually from N to Q using a Q5 Site-Directed Mutagenesis Kit (NEB). The full-length or C19 truncated fragments were amplified and inserted into plvx-puro vector. Western blot analysis of protein cleavage and pseudovirus packaging was performed as described.

### Western blot analysis

Unless noted, proteins were separated by SDS-PAGE on 4–20% Bis-Tris gels and transferred to PVDF membranes. Western blotting was performed using a Pierce Fast Western Blot Kit (Thermo Scientific) according to the manufacturer’s instructions. SuperSignal West Femto Maximum Sensitivity Substrate (Thermo Scientific) was used for chemiluminescence detection where needed. The following antibodies were used: anti-S2 antibody (which detects the S2 subunit of SARS-CoV-2 and SARS-CoV) was from Novus (NB100-56578); anti-FLAG antibody was from Sigma (F1804); anti-gp120 was from NIH-ARP (clone 16H3, ARP-12559); anti-RPN2 was from ABClonal (A8352); and anti-HA (51064-2-AP), anti-MYC (16286-1-AP), anti-GBP5 (13220-1-AP), anti-RPN1 (12894-1-AP), anti-DDOST(14916-1-AP), anti-STT3B (15323-1-AP), anti-STT3A (12034-1-AP), HRP-conjugated anti-α-tubulin (HRP-66031), and HRP-conjugated anti-GAPDH (HRP-60004) antibodies were all purchased from Proteintech. The signal intensity of western blot bands was quantified by ImageJ (https://imagej.nih.gov/ij/) and indicated under the figure panels.

### Immunofluorescence microscopy

Transfected 293T cells were fixed with 4% PFA, permeabilized with 0.1% Triton X-100 in PBS, blocked with 5% BSA in PBS for 1 h, and then incubated with primary antibodies at 4°C overnight. After washing, the cells were incubated with secondary antibodies for 1 h. Hoechst 33342 was used to stain the nuclei. Images were acquired using a Leica SP8 confocal microscope. Anti-calnexin (C4731) was from Sigma-Aldrich, anti-TGN46 (NBP149643) was from Novus Biological, and goat anti-rabbit IgG 594 (A-11012) was from Invitrogen.

### Statistical analysis

Data are presented as the mean ± standard deviation unless indicated and were analyzed using Student’s t-test (paired, two-sided) with Prism 9 (GraphPad). The n value represents the number of independent experiments. A p value of <0.05 was considered statistically significant.

## ACKNOWLEDGEMENTS

We thank members of the Rana lab for helpful discussions and advice. The following reagent was deposited by the Centers for Disease Control and Prevention and obtained through BEI Resources, NIAID, NIH: SARS-Related Coronavirus 2, Isolate USA-WA1/2020, NR-52281. We thank the UC San Diego Center for Advanced Laboratory Medicine Clinical Microbiology and Virology Lab and UC San Diego EXCITE for providing clinical samples for viral isolation and SARS-CoV-2 genome sequencing. This work was supported by a Career Award for Medical Scientists from the Burroughs Wellcome Fund USA and a grant from the National Institutes of Health USA (K08 AI130381) to AFC, and in part by grants from the National Institutes of Health (CA177322, DA039562, DA046171, and AI125103).

## AUTHOR CONTRIBUTIONS

SW and WL designed and performed experiments, analyzed the data, and wrote the manuscript; LW, SKT, WB, LW, AEC, QZ, LZ performed experiments and analyzed the data; NL and HH analyzed the data; AFC, advice in experimental plans and data analysis; TMR, conceived the overall project and participated in experimental design, data analyses, interpretations, and manuscript writing.

