## Supplementary figures and images for "Interferon-Inducible Guanylate-Binding Protein 5 Inhibits Replication of Multiple Viruses by Binding to the Oligosaccharyltransferase Complex and Inhibiting Glycoprotein Maturation"

### Supplementary Figures 1 to 5

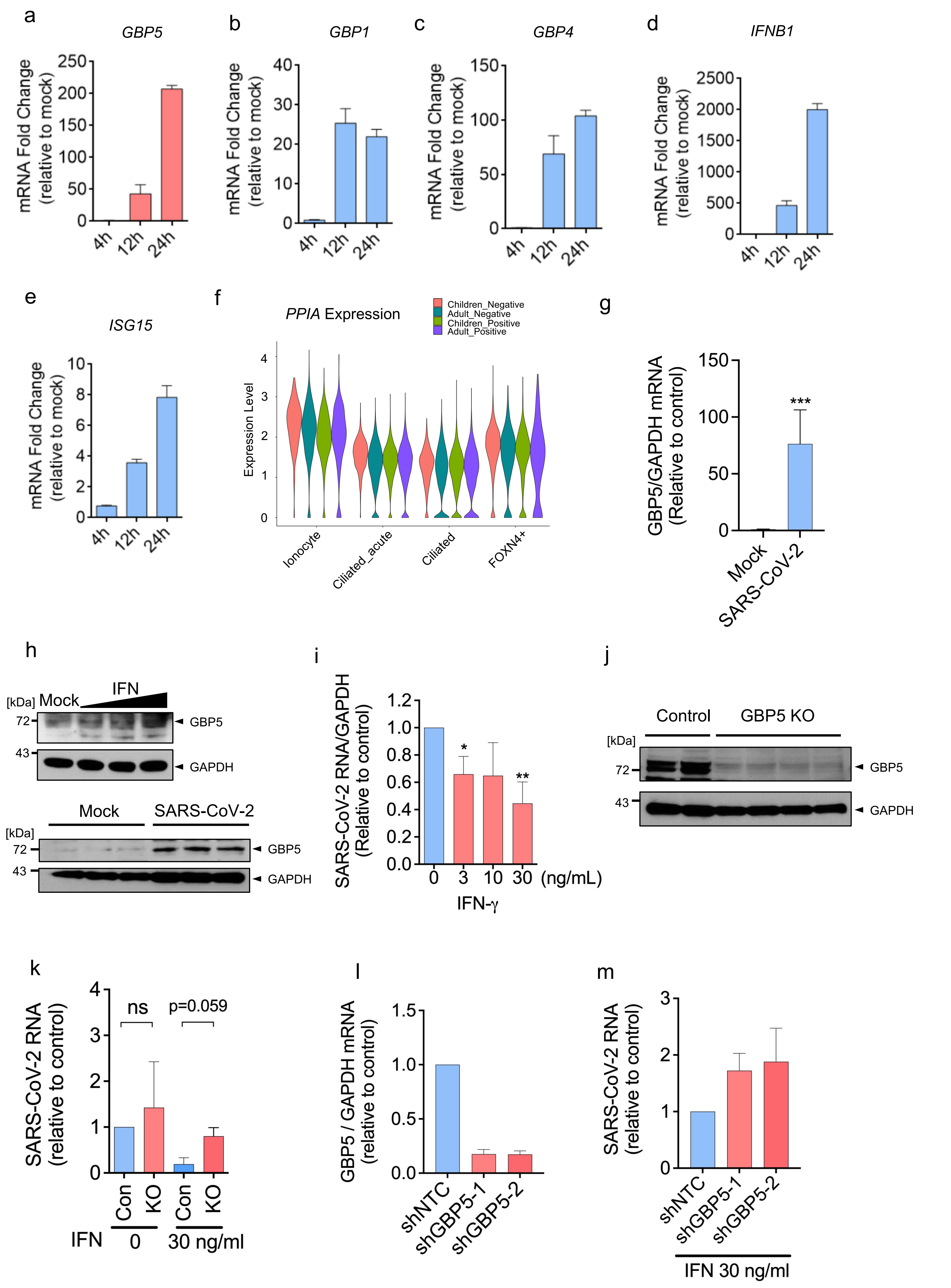

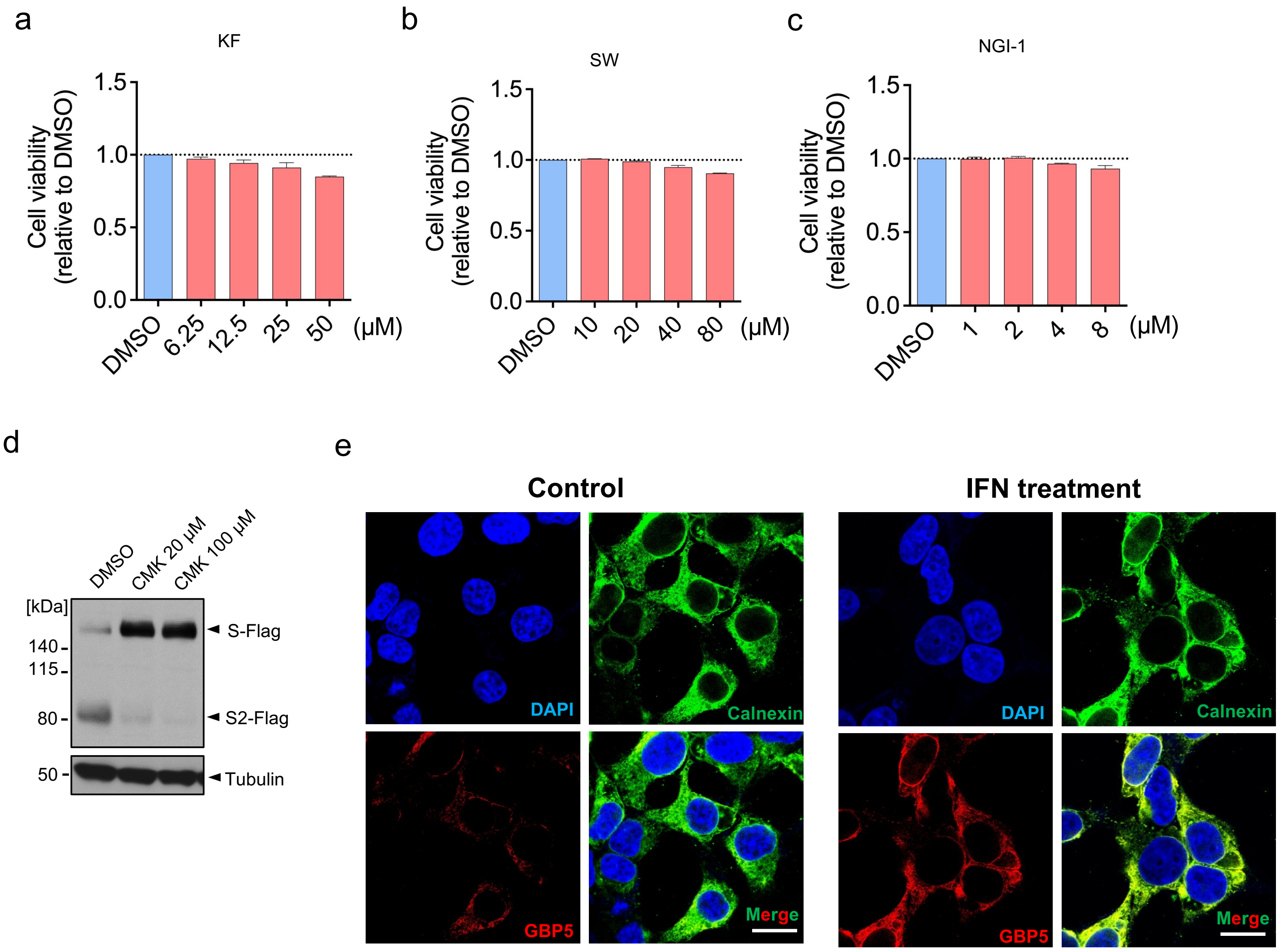

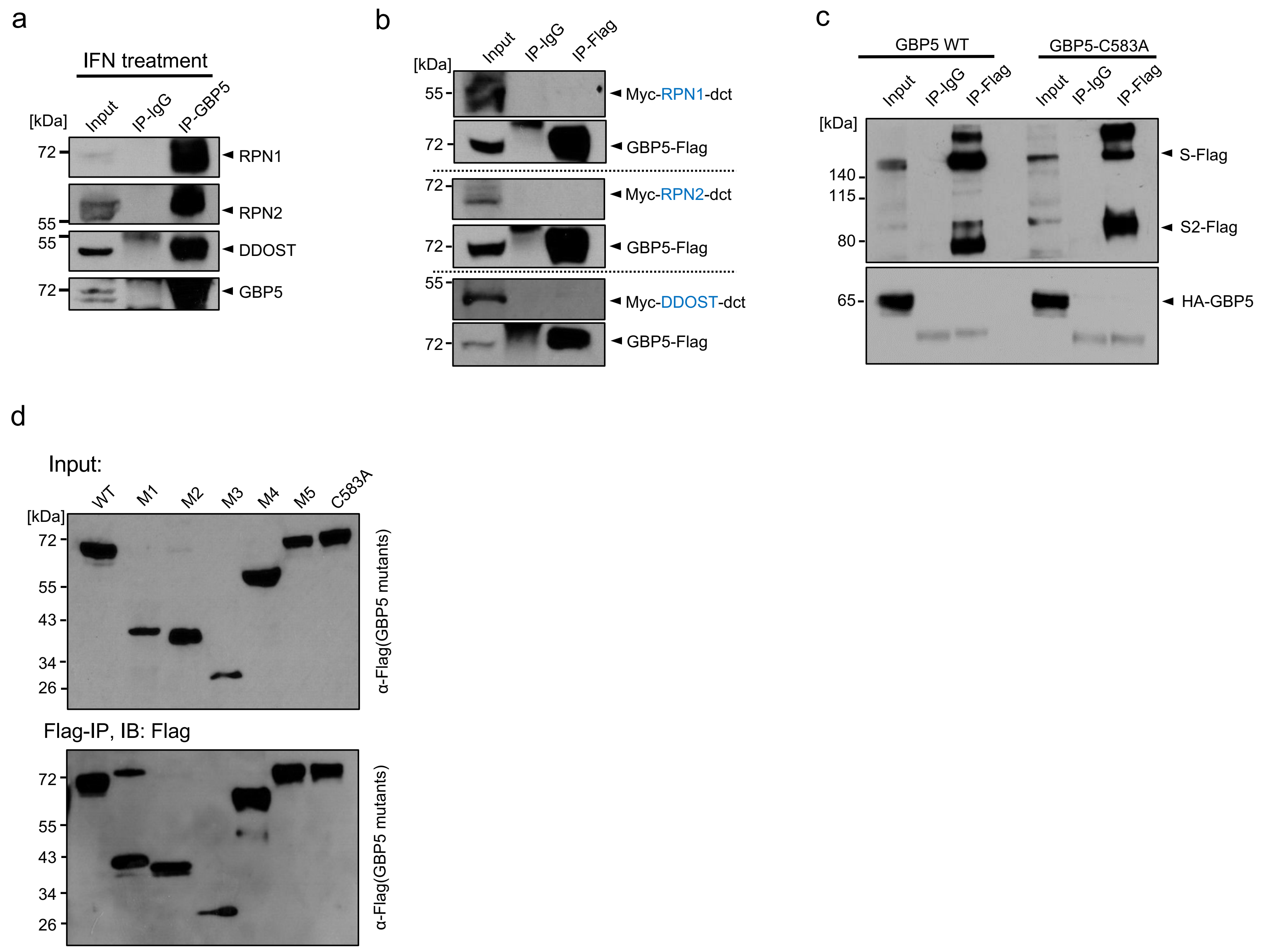

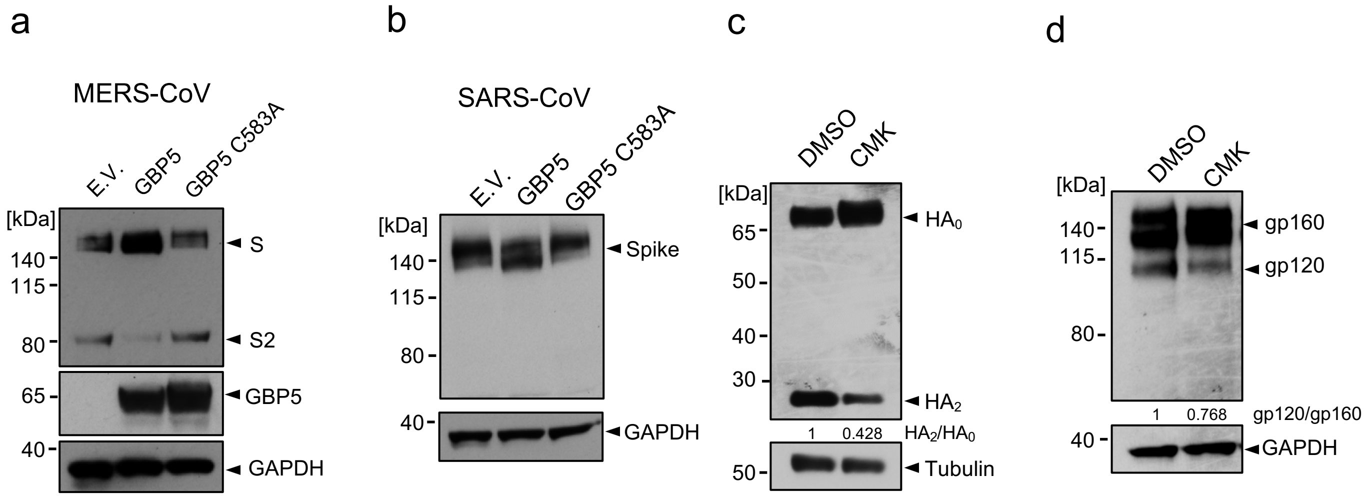

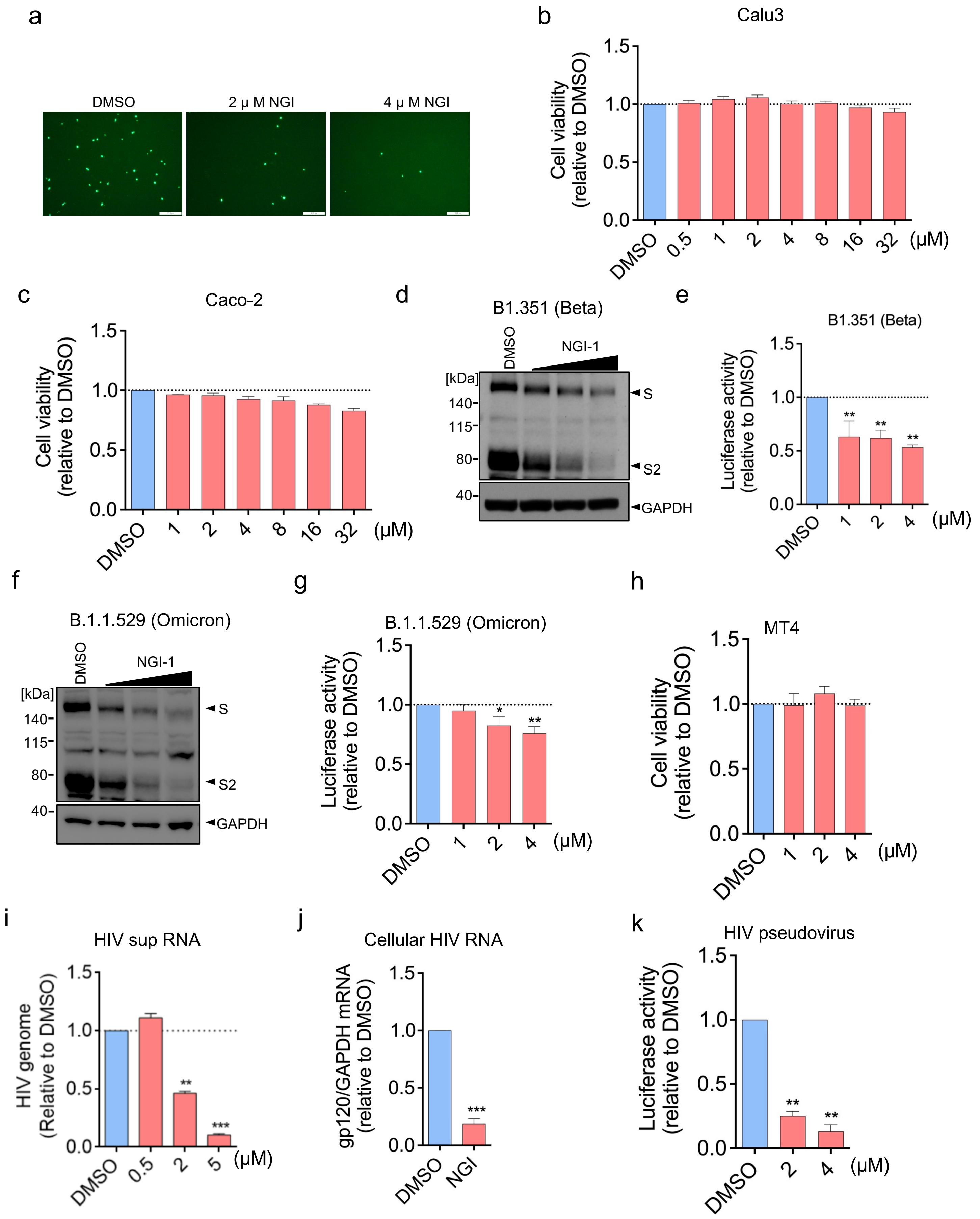
